# Long-term repair of porcine articular cartilage using cryopreservable, clinically compatible human embryonic stem cell-derived chondrocytes

**DOI:** 10.1101/2021.05.27.446024

**Authors:** Frank A. Petrigliano, Nancy Q. Liu, Siyoung Lee, Jade Tassey, Arijita Sarkar, Yucheng Lin, Liangliang Li, Yifan Yu, Dawei Geng, Jiankang Zhang, Ruzanna Shkhyan, Jacob Bogdanov, Ben Van Handel, Gabriel B. Ferguson, Youngjoo Lee, Svenja Hinderer, Kuo-Chang Tseng, Aaron Kavanaugh, J. Gage Crump, April D. Pyle, Katja Schenke-Layland, Fabrizio Billi, Liming Wang, Jay Lieberman, Mark Hurtig, Denis Evseenko

**Author notes:** Correspondence to: Denis Evseenko MD, PhD. Associate Professor of Orthopaedic Surgery, Stem Cell Research and Regenerative Medicine. USC, Los Angeles, CA, 90033, USA. 1450 Biggy St. NRT4509, Los Angeles, CA 90033. These authors contributed equally to this work (Frank A. Petrigliano, Nancy Q. Liu, Siyoung Lee, Jade Tassey).

## Abstract

Osteoarthritis (OA) impacts hundreds of millions of people worldwide, with those affected incurring significant physical and financial burdens. Injuries such as focal defects to the articular surface are a major contributing risk factor for the development of OA. Current cartilage repair strategies are moderately effective at reducing pain but often replace damaged tissue with biomechanically inferior fibrocartilage. Here we describe the development, transcriptomic ontogenetic characterization and quality assessment at the single cell level, as well as the scaled manufacturing of an allogeneic human pluripotent stem cell-derived articular chondrocyte formulation that exhibits long-term functional repair of porcine articular cartilage. These results define a new potential clinical paradigm for articular cartilage repair and mitigation of the associated risk of OA.

---

Cell therapy has been used successfully in the clinic for more than 50 years in the form of hematopoietic stem cell transplantation^1^. This pioneering work illuminated the need for HLA-matched donors due to graft versus host disease (GVHD) encountered during allogenic transplants, in which donor lymphocytes reacted against host tissues. In the case of allogenic solid organ transplantation, immunosuppression of the host is often required for extended periods^2^. These aforementioned limitations in the availability and compatibility of donor tissue have prompted the search for other solutions which now potentially include human embryonic stem cell- (hESC) and induced pluripotent stem cell-derived (iPSC) cells and tissues^3,4^. The field of PSC-based regenerative medicine has advanced quickly as both iPSC- and ESC-derived cell therapies are in clinical trials^5^, with transplants into immunoprivileged sites such as the eye leading the way.

In the orthopaedic field, reparative therapy for articular cartilage defects has classically relied on endogenous cells via the microfracture technique. In this procedure, channels are created through the subchondral plate into the bone marrow to allow mesenchymal stromal cells (MSCs) with chondrogenic potential to enter the defect and generate neocartilage^6^. The reparative fibrocartilage produced following microfracture is often biomechanically inferior to the surrounding hyaline articular cartilage, as MSCs are undergoing chondrogenesis in an inflammatory microenvironment^7^ and in the absence of inductive cues such as BMP-2^8^; augmented microfracture techniques that address some of these potential limitations are being actively explored^9^. More recently, autologous chondrocyte implantation^10^ (ACI) and variations thereof including MACI^11^ (matrix-associated ACI) that rely on expansion and reimplantation of chondrocytes from the patient, have been adopted. The long-term results of ACI-based procedures appear superior to microfracture, with reduced graft failure and improved patient-reported outcomes^12,13^. Despite overall satisfactory outcomes, many patients still experience complications including graft integration failure, inferior quality of neocartilage (hyaline vs. fibrocartilage), donor site morbidity, osteoarthritis^14^ and chronic pain. Additionally, these approaches require two staged surgical procedures to perform the actual repair, increasing the overall cost and logistic complexity. This has fueled the search for allogeneic sources that may provide more cells with superior chondrogenic capacity without immunocompatibility issues or the need for multiple surgeries including juvenile cartilage^15^ and PSCs.

Generation of articular chondrocytes from PSCs has been challenging as most chondrogenic cells during development are fated to undergo hypertrophy and endochondral ossification rather than adopt an articular chondrocyte identity^16^. We^17,18^ and others^19-22^ have generated articular-like chondrocytes from human pluripotent stem cells; we have subsequently shown that stable articular chondrocytes produced from GFP^+^ PSCs can engraft, integrate into and repair osteochondral defects in small animal models^18^. Moreover, these human cells produce all layers of hyaline cartilage after 4 weeks *in vivo*, including a PRG4^+^ superficial zone^18^. However, production scaling and assessment of long-term, clinically relevant functionality has so far limited the development of these protocols. The Yucatan minipig presents an excellent model for pre-clinical assessment of potential orthopaedic therapies due to structural similarities, comparable thickness of articular cartilage and the ability to create defects of substantial volume^23-25^; in addition, their size allows for cost-efficient care and observation for extended periods of time. Here we present data demonstrating long-term functional repair of porcine full-thickness articular cartilage defects with hyaline-like cartilage by scalable production of clinical grade hESC-derived immature articular chondrocytes.

## Results

### Scaled production and formulation optimization of hESC-derived chondrocytes

The main purpose of this study was to assess the long-term therapeutic potential of hESC-derived chondrocytes in a porcine model of focal articular cartilage injury. We have previously defined a protocol for the generation of articular cartilage-like chondrocytes from human PSCs^17^. Cells generated using this technique are immature based on their transcriptional signature and expression of immature chondrocyte markers but can mature upon implantation *in vivo*, evidencing appropriate of superficial zone markers and lack of hypertrophy^18^. For this study, we have adapted our previous protocols^17,18^ initially developed for H1 and H9 lines, to utilize the research grade hESC line ESI-017^26^. This specific line was selected because a cGMP version of this line is fully compliant with all current FDA regulations and can be advanced into human clinical trials without any regulatory restrictions.

In order to generate sufficient numbers of clinical grade hESC-derived chondrocytes for cartilage defect repair, hESCs were first expanded and induced into mesodermal differentiation (d1-7) followed by chondrogenic differentiation (d7-11; Figure 1a). At d11, mesodermal skeletal progenitors were isolated using MACS to deplete for epithelial (undifferentiated and epidermal, EpCAM/CD326^+^) and cardiovascular mesodermal (KDR/CD309^+^)^27^ cells (Figure 1a).

**Figure 1:**
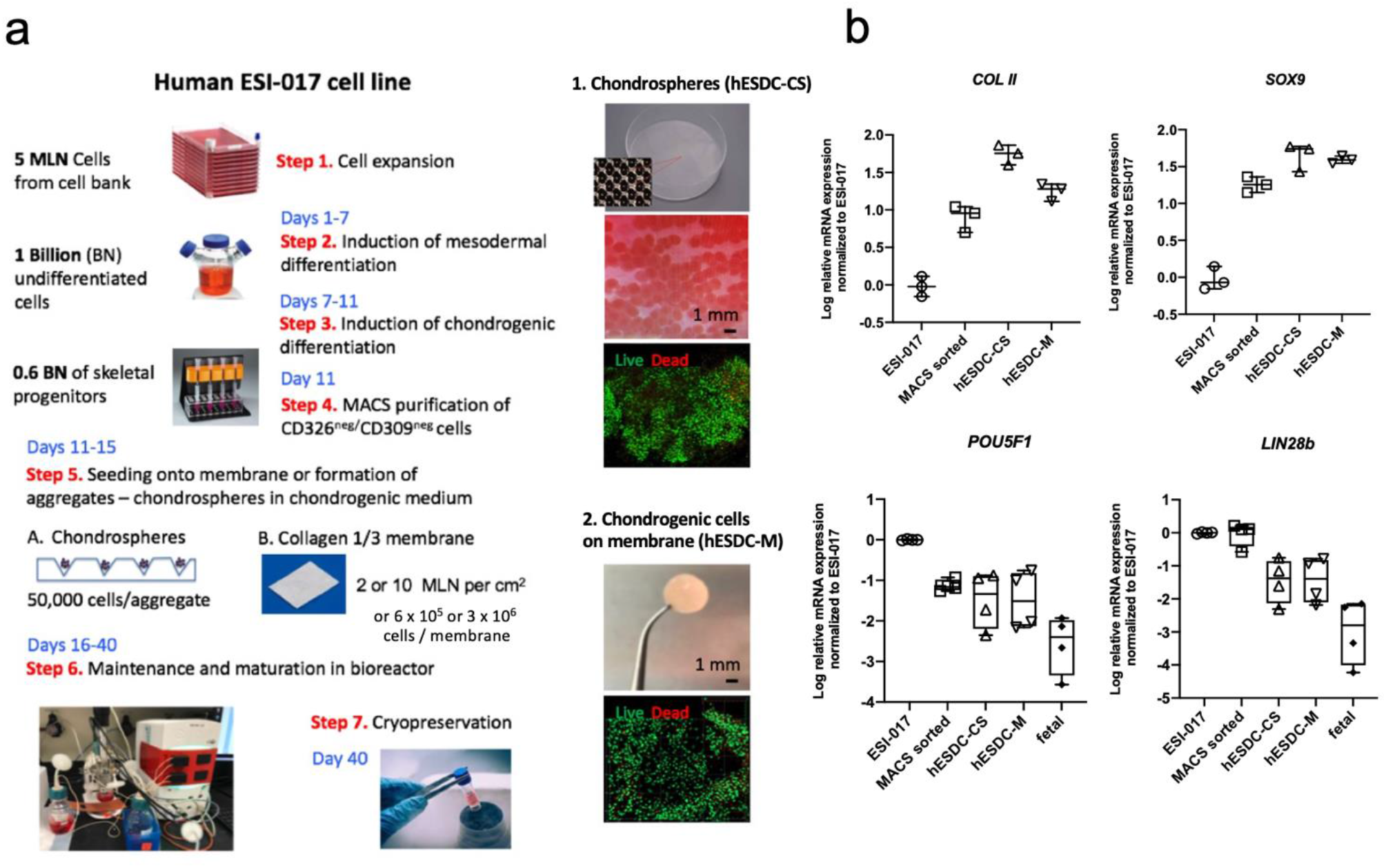
Scale up and formulation of cGMP-grade hES-derived chondrocytes. (a) Schematic depicting the large-scale production of chondrocytes in 2 different formulations from ESI-017 cells. Pre-chondrocytes were seeded onto clinically-used porcine collagen I/III membranes (M) or aggregated to generate chondrospheres (CS). Cells were expanded and then cryopreserved under optimal conditions described in Supplementary Figure S1. (b) qPCR for chondrogenic and pluripotent genes at different stages of *in vitro* differentiation or *in vivo* fetal ontogeny (14-17 weeks). n=4 different batches or biological replicates (fetal); data presented as box and whisker plots showing all points.

During the optimization stage we produced and tested 2 different formulations of hESC-derived chondrocytes: chondrospheres (CS) and chondrocytes integrated onto a collagen I/III membrane previously approved for clinical use^28^ (Cartimaix; Matricel). For chondrosphere production, we used a commercially available low attachment plate with a patterned floor designed to generate chondrospheres of uniform size and quality from d11 MACS-purified chondrogenic cells (Figure 1a). In parallel, collagen membranes were sized to 6 mm (0.28 cm^2^) with a biopsy punch and seeded with purified skeletal progenitors isolated after MACS and transferred aseptically into the bioreactor. We then used a continuous perfusion bioreactor system (Figure 1a) for expansion and chondrocyte maturation for an additional 25 days to provide a stable microenvironment with precise control of gases, nutrients and physical parameters such as shear stress. We confirmed that ESI-017 cells responded to our established protocol by upregulating chondrogenic and downregulating pluripotency genes during the course of the manufacturing process (Figure 1b.

We then tested cryopreservation media (Figure S1) for each of these formulations to support the development of a universal, off-the-shelf potential therapy for articular cartilage defects. After optimization of cryopreservation, both chondrospheres and membranes were revived with >70% viability of cells (Figure S1). These batches routinely yielded dozens of chondrospheres containing ∼5 × 10^4^ cells, or low and high dose membranes containing 6 × 10^5^ or 3 × 10^6^ immature chondrocytes each, respectively.

We then compared the ability of two doses of each formulation to support short-term repair of a focal defect in pig articular cartilage (Figure S2). Six millimeter full-thickness cartilage defects were created to the depth of the subchondral plate in the femoral condyle and low and high doses of hESC-derived chondrospheres or the same cell numbers of chondrocytes on membranes were implanted with fibrin glue (Figure S2a); empty membranes or glue alone were used as controls. One month later, injured areas were assessed for evidence of tissue integration, matrix production and fibrosis; detailed biomechanical mapping was also conducted to determine biomechanical characteristics of the healing defects. These results (Figures S2,3) clearly demonstrated that a high dose of chondrocytes embedded in collagen membranes was better maintained in the injury site and supported superior functional repair without eliciting a heightened immune response (Figure S4^29^). Based on these data, we proceeded with membrane-bound hESC-derived chondrocytes (hESDC-M) for further characterization and functional testing.

### Characterization and developmental status of hESDC-M

We have previously shown chondrocytes derived from hESCs using our protocol may lie between fetal and adult cells on the developmental timeline^18^. These experiments were previously conducted on cells isolated from chondrospheres using bulk RNA-seq. To better understand the heterogeneity present in hESDC-M, we performed single cell RNA-seq (scRNA-seq) on cells isolated from two different batch production runs and compared them to undifferentiated ESI-017 cells (Figure 2). Importantly, few to none of the cells demonstrated expression of pluripotency genes (*POU5F1, LIN28A or ZFP42*; Figure S5a), suggesting limited potential for generation of teratomas upon transplantation *in vivo*. Both production runs contained similar cell types as shown by objective clustering (Figure 2a), with the absolute majority of cells present being positive for chondrogenic markers including *COL2A1, HAPLN1*, and *SOX5/6/9*, (Figures 2b-d). This was affirmed by gene ontology analysis of genes enriched in the hESDC-M (FDR <0.05, >2-fold change) as demonstrated by significant over-representation of genes related to cartilage development, matrix production and lineage commitment (Figure 2e). Objective clustering of hESDC-M defined 4 subtypes of cells present on the membranes, with clusters 1 and 2 being most enriched for *COL2A1* and *PRG4* (Figure 2f-h). To define how these subtypes of hESDC-Ms compare with different stages of human ontogeny and human bone marrow MSCs cultured on membranes (hBM-MSC-M), we performed scRNA-seq at multiple stages of human chondrogenic ontogeny and on hBM-MSC-M (Figures S5,6). These data defined cluster 1 as the most mature, with highly significant overlap with genes enriched in fetal chondrocytes vs. embryonic chondroprogenitors, including genes encoding ECM proteins such as *COL2A1, PRG4* and *ACAN* (Figures 2i,j; S6). Clusters 2 and 3 were more closely related to chondroprogenitors, showing enrichment for genes involved in the generation of chondrogenic condensations and primitive mesoderm including *TWIST1*^*30*^, *NCAM1* and *CDH2*^*31*^ (Figures 2k, l; S6). These data were confirmed by comparison to our previous bulk sequencing data generated at these stages^17,18^ (Figure S6e). Cluster 4 contained cells expressing *COL2A1* and neural markers including *PAX6* and *MITF* (Figure 2m), similar to a population recently described to be present during hIPSC chondrogenesis^32^. Conversely, hBM-MSC-M cultured under identical conditions to hESDC-M expressed genes associated with terminal chondrogenesis including *COL10A1* and *SPP1* (Figure S5c,d). These data reveal that although the cells present on the membrane are heterogenous, the production process is reproducible and generates mostly immature chondrogenic and chondroinductive cells.

**Figure 2:**
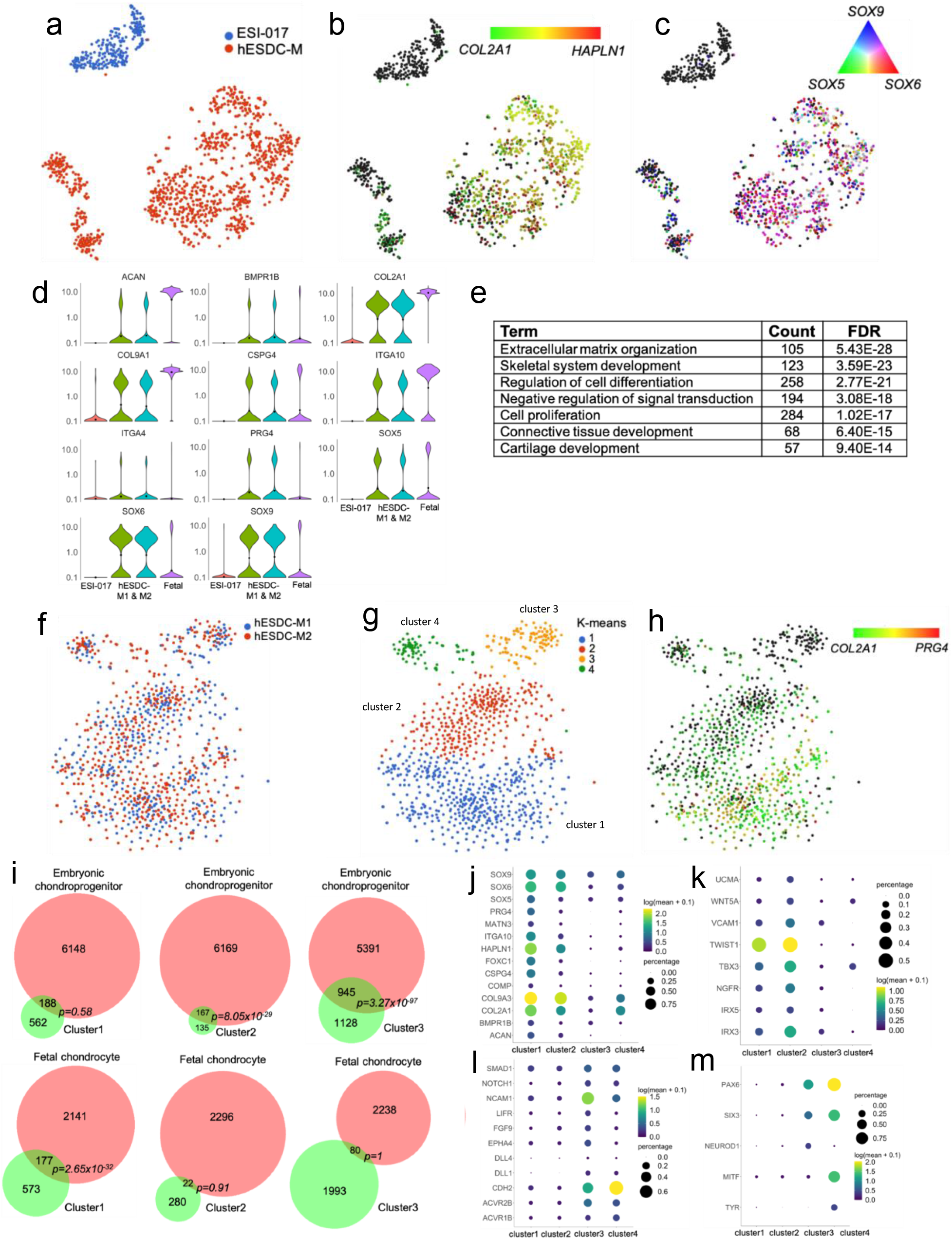
Transcriptional profiling of membrane embedded hESDC-M. (a) t-SNE plot of single cell sequencing data generated from ESI-017 cells (n=1, blue) or hES-derived chondrocytes digested from membranes at d40 of differentiation (n=2; cells pooled in red). (b,c) t-SNE plots depicting expression of indicated genes at single cell resolution. (d) Violin plots for gene expression of selected chondrogenic genes; fetal chondrocyte expression data are shown for reference. (e) Selected gene ontology (GO) categories enriched in hESDC-M vs. ESI-017 cells based on genes with FDR <0.05, >2-fold change. (f) Re-clustering, (g) k-means clustering of hESDC-M and (h) *PRG4* and *COL2A1* expression levels. (i) Venn diagrams demonstrating overlap of biomarker genes of the indicated cluster with (top) genes enriched in embryonic chondroprogenitors vs. fetal chondrocytes analyzed by scRNA-Seq or (bottom) vice versa. (i-m) Expression of selected and biomarker genes in each cluster of hESDC-M.

### hESDC-M support long-term repair of articular cartilage in pigs

In order to assess the therapeutic potential of hESCD-M, we designed a long-term clinically relevant experiment in which either membranes alone or membranes with cells were implanted into pig articular cartilage defects and assessed 6 months later. Pigs in each group (n=5) had 2, 6 mm full-thickness cartilage defects created in their femoral condyles within the load bearing areas and were treated with either membranes alone or hESDC-M; two animals were used as sham controls with no cartilage defects generated. Cell implants were thawed and washed in fresh X-Vivo media approximately 1 hour prior to implantation, applied to the defects, and were fixed in place with fibrin glue. After 6 months, pigs were euthanized and cartilage assayed using morphological, histological and biomechanical methods. Visually, defects from all pigs transplanted with cells uniformly evidenced substantially less degeneration in and around the injury site (Figure 3a). Sham operated animals showed no noticeable morphological or biomechanical differences with non-operated knees (data not shown). At the microscopic level, defects treated with cells contained neocartilage with more proteoglycan deposition and better integration of the new cartilage tissue with the non-injured surrounding matrix (Figures 3b, S4e). To quantify the extent of regeneration provided by cells, all 10 defects per group were scored by 2 blinded observers using the International Cartilage Repair Society (ICRS) II histological assessment system^33^ (Figure 3c). This scoring system grades 14 criteria relevant to cartilage repair and provides a comprehensive view of the utility of potential treatment. Scores from each observer for each defect were averaged to provide a composite for each criterion. These data showed significantly better outcomes in most categories for defects treated with hESDC-M (Figure 3c). This was confirmed by synovitis scoring and staining for inflammatory infiltrates, which showed no difference between animals implanted with empty membranes vs. those receiving hESDC-M, further supporting the low immunogenicity of hESDC-M (Figure S4). Moreover, morphological signs of synovitis were more prominent in pigs treated with membranes only, likely reflecting progression of degenerative joint disease in this group; however, this difference did not reach statistically significant values (Figure S4d). Histological analysis of the subchondral bone showed no major differences between the groups (data not shown).

**Figure 3:**
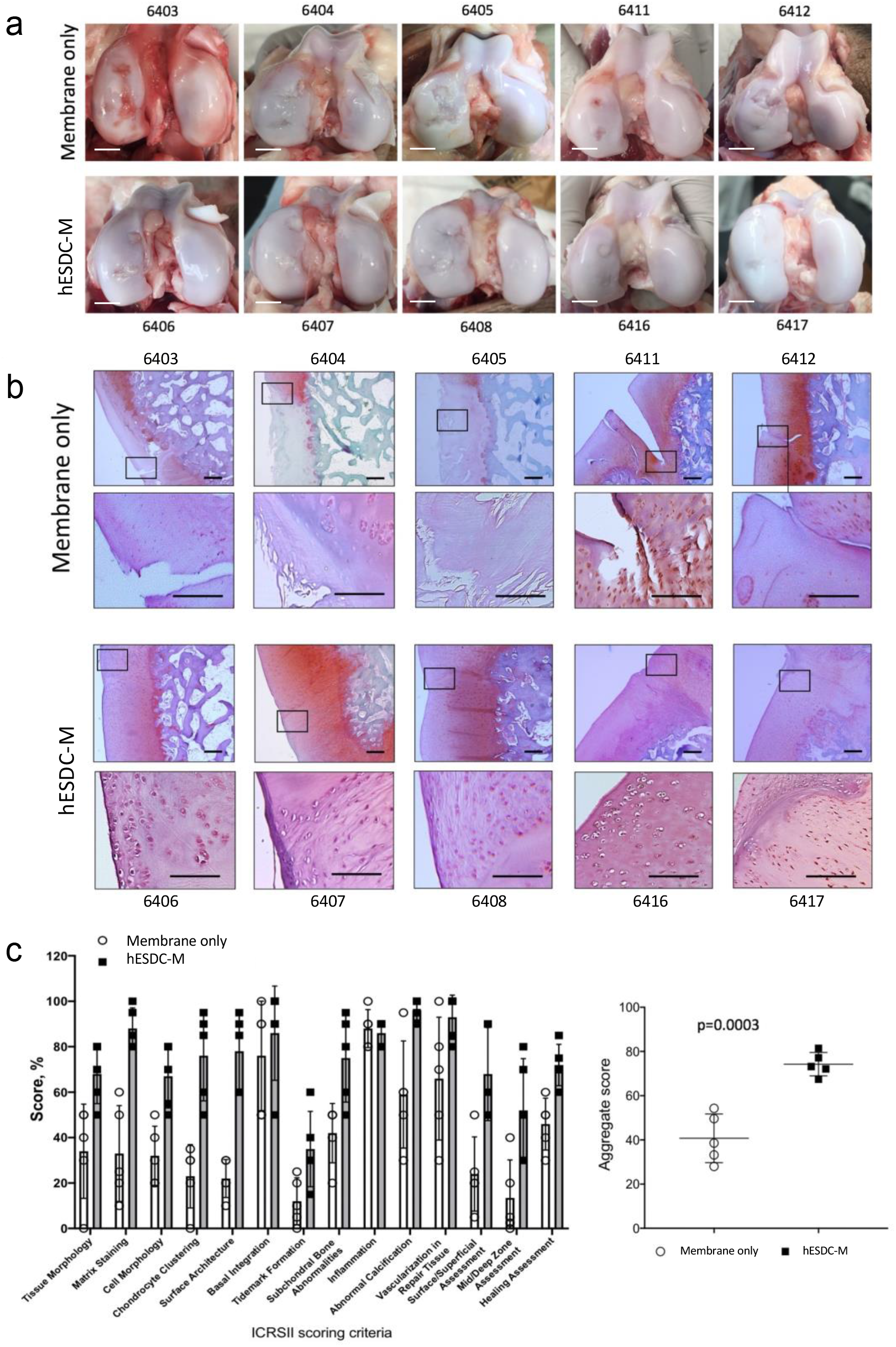
Focal articular cartilage defects treated with hESDC-M show improved repair at 6 months. (a) Gross visual appearance of all 10 defects created in the femoral condyle of control (membrane alone, top row) or treated (hESDC-M, bottom row) Yucatan minipig knees after 6 months. Scale bar = 10 mm. (b) Safranin O/Fast Green staining of the interface between the graft and endogenous tissue or the defect itself (boxes); where the boxed regions are shown at higher magnification below. Scale bar = 100 μm. (c) Histological scoring of sections from control and treated femoral condyles for the 14 parameters comprising the ICRS II cartilage repair scoring system (left); each point represents the average of both defects per animal. (Right) Aggregate score of all 14 parameters over the 10 defects scored. Identifiers above or under images represent each animal. p-value was calculated using unpaired Student’s t-test; data presented as mean ± SD.

At the molecular level, cartilage in the defects treated with cells more closely resembled the surrounding tissue. In membrane only defects, much of the new tissue was inferior as evidenced by collagen 1 and collagen X staining coupled with substantially reduced proteoglycan content (Figure 4a). In contrast, defects transplanted with hESDC-M evidenced appropriate stratification of the neocartilage as demonstrated by superficial production of lubricin (PRG4) and localization of SOX9^+^ cells (Figure 4a). Moreover, substantial production of collagen II in the transitional zone was primarily observed in defects treated with cells. Notably, even after 6 months following implantation, small clusters of Ku80^+^ human cells^34^ were identifiable in all treated animals (Figure 4b). With the safety profile of hES-derived chondrocytes in mind, we explored the biodistribution of human cells by using a sensitive PCR-based assay to detect human telomerase (*TERT*) in cartilage, synovium, peripheral blood and major organs (Figure 4c,d). While the presence of human cells in the repaired articular cartilage was confirmed and represented roughly 4% of total cells (Figure 4b), levels of human DNA in all other tissues analyzed, including synovium, was below the threshold of detection, indicating that transplanted cells do not leave the defect following implantation after 6 months.

**Figure 4:**
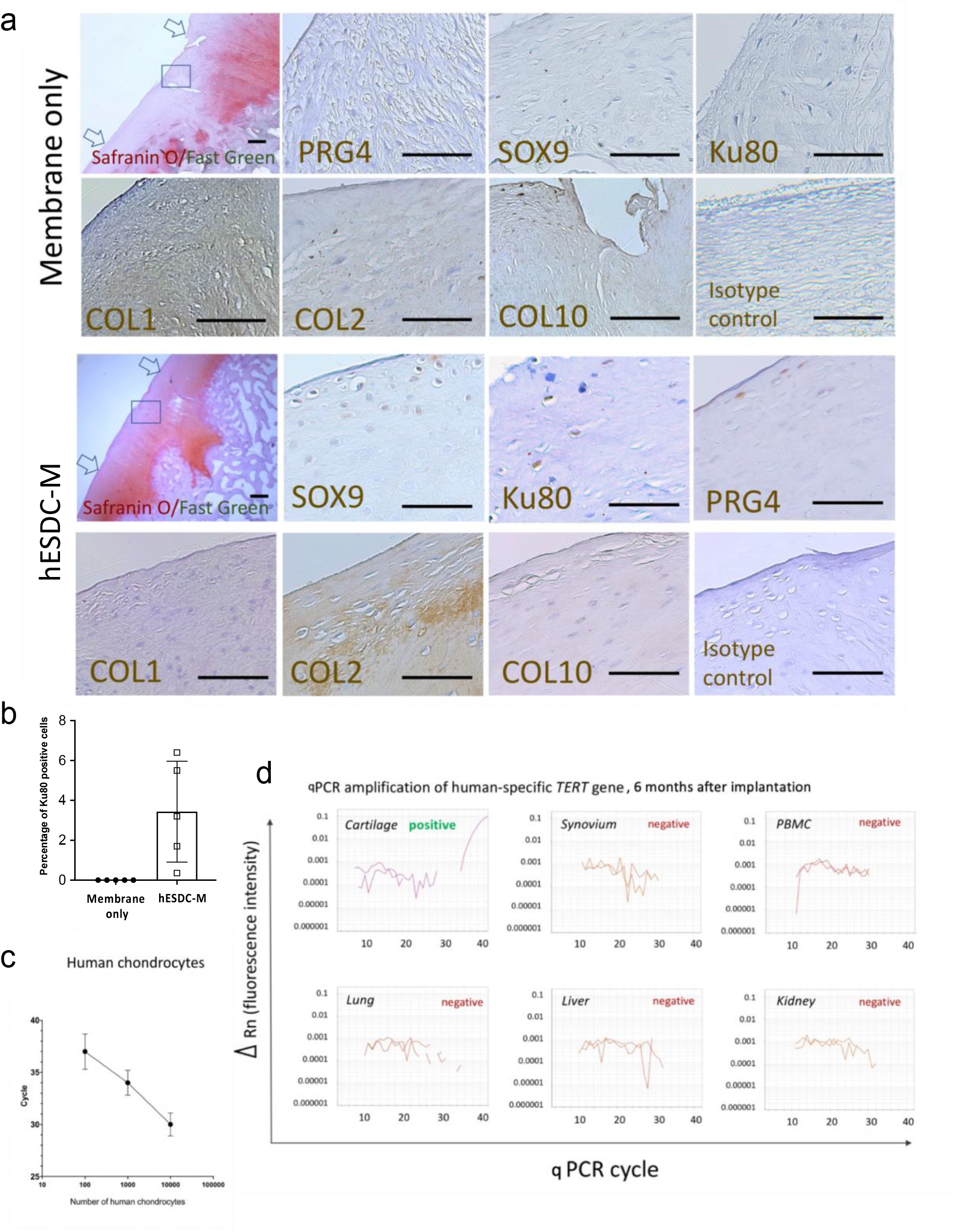
hESDC-M treated defects evidence superior repair and contain both human and pig cells at 6 months. (a) Immunohistochemical staining of the full defect (indicated by arrows) for Safranin O/Fast Green to assess glycosaminoglycans for control (membrane only) and treated (hESDC-M) animals. Representative images of immunohistochemical staining of the boxed area for human-specific antigen Ku80 and zonal markers of articular cartilage for both control and treated femoral condyles are shown; scale bar = 200 μm. (b) Quantification of Ku80+ cells (mean ± SD of 5 biological replicates). (c) qPCR analysis of human *TERT* gene. Standard curve constructed with human chondrocyte genomic DNA allowed reliable detection of as few as 100 human cells (mean ± SD of 3 biological replicates). (d) Genomic DNA extracted from the indicated tissues was analyzed for the human *TERT* gene. Representative amplification plots are shown; human cells were detected in all defects of animals treated with hESDC-M. PBMCs = peripheral blood mononuclear cells.

As a measure of functional repair, we evaluated the biomechanical properties of the neocartilage in control and cell-treated defects (Figure 5). The surfaces of both femoral condyles were biomechanically mapped in all 10 pigs to assess cartilage stiffness and thickness (Figure 5a). The instantaneous modulus, a measure of compressibility of cartilage, was higher in most defects treated with cells and more similar to native cartilage (Figure 5b). As expected from the histology, cartilage in and around the defects treated with cells was thicker than in defects treated with membranes alone (Figure 5a,c). Finally, the biomechanical properties of all repaired defects in each group were merged into a composite score; although animals treated with cells did not achieve full restoration of biomechanics, they were significantly improved as compared to defects treated with membranes alone (Figure 5b,c). These results indicate that hESDC-M promote the formation of hyaline cartilage that is biomechanically similar to native tissue for at least 6 months after implantation.

**Figure 5:**
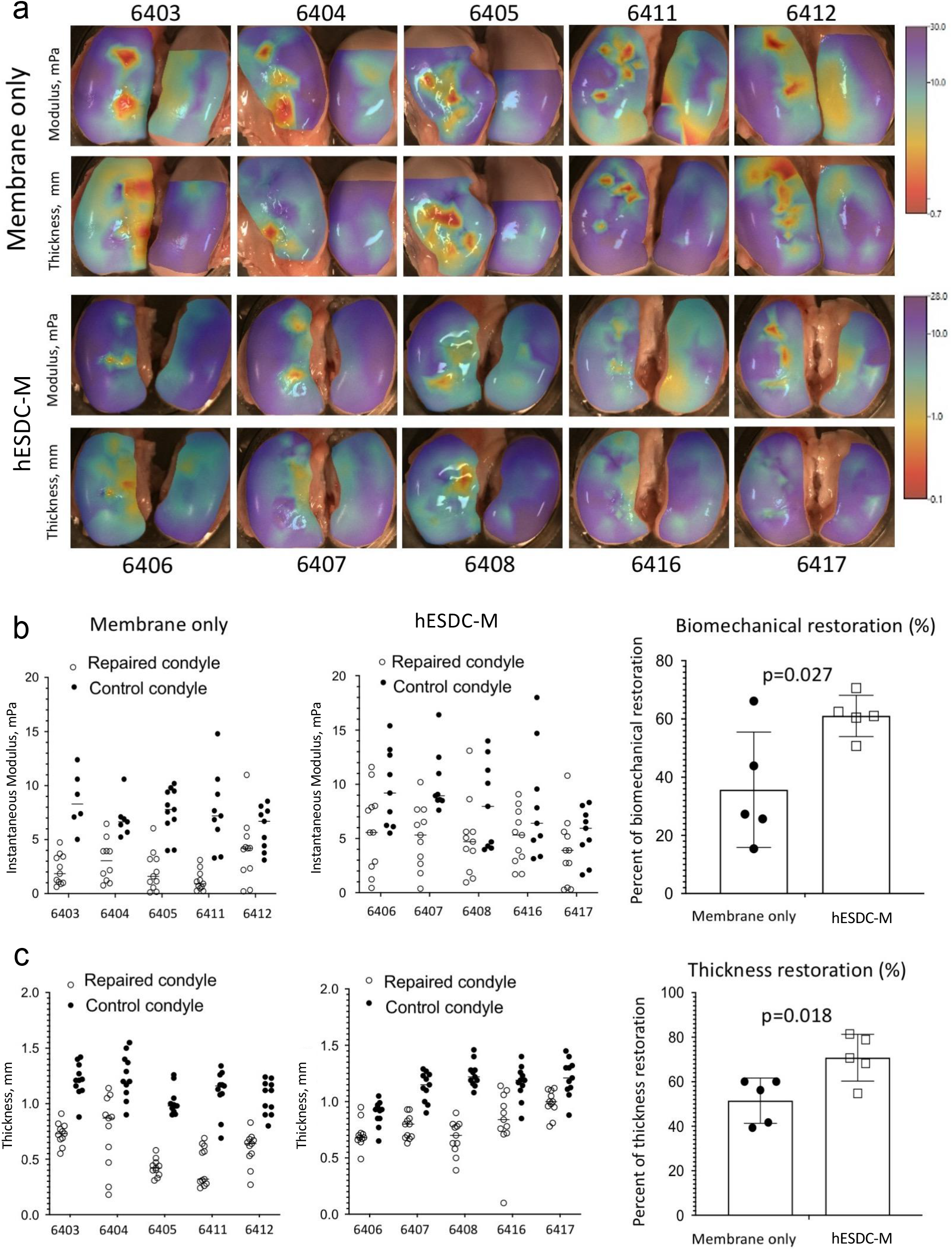
hESDC-M elicit biomechanically superior articular cartilage repair long term in porcine knees at 6 months. (a) Heat maps depicting scanning indentation and thickness of femoral condyles generated using Mach-1 bioindentor; scale bars for instantaneous modulus (top rows) and thickness (bottom rows) are shown on the right. (b) Data points for instantaneous modulus or (c) thickness measurements for each defect (2 per condyle) were merged into one aggregate measure and compared to the same area of the uninjured condyle; values for both defects per animal were averaged and the plotted as controls (membrane only) vs. treated (hESDC-M) as a function of average uninjured condyle measurements to calculate percent restoration (right). Identifiers represent individual animals (n=5). p-value was calculated using unpaired Student’s t-test; data presented as mean ± SD.

### hESDC-M secrete chondroinductive factors that induce chondrogenesis from porcine MSCs

Given that the majority of hyaline-like neocartilage was contributed by pig cells, we assessed the paracrine effects of hESDC-M (Figure 6). We utilized a methylcellulose (MC)-based culture method to assess both clonality at a single cell level and capacity of pig MSCs to undergo chondrogenesis *in vitro* (Figure 6a). Growth factors of the TGF-β^35^, FGF^36^ and BMP^37^ families are known to be chondroinductive during development and following injury. Upon this basis, three selected growth factors (FGF-2, BMP-2, and TGF-β1, 3GFs) produced by hESDC-M (Figure S7) were added to the MC-based media, showing that a combination of all 3 growth factors yielded the most clones (Figure 6b); notably, BMP-2 was not secreted by hBM-MSCs (Figure S7e). These colonies produced proteoglycans and other chondrogenic markers comparable to native articular chondrocytes cultured in the same method (Figure 6c, S8e). Chondrogenic gene expression similar to that of native pig articular chondrocytes was evident when comparing MSCs cultured in MC with 3GFs, with substantial induction of chondrogenic genes versus starting MSCs (Figure 6d). Moreover, culture in MC + 3GFs yielded chondrogenic gene expression similar to micromass culture of pig MSCs + 3GFs, the standard method for generating chondrocytes from MSCs *in vitro*. To assess whether paracrine factors produced by hESDC-M could promote chondrogenesis from pig MSCs, we employed a co-culture system with Transwell inserts and MC (Figure 6e). After 4 weeks of co-culture, clonality and chondrogenesis showed the same trends as MSCs cultured with all 3GFs (Figure 6f,g); expression of collagen X was undetectable. Importantly, hESDC-M also supported clonal chondrogenesis from pig chondrocytes (Figure S8), suggesting two possible cellular sources for neocartilage following hESDC-M implantation. These data indicate secreted proteins such as BMP-2 produced by hESDC-M can promote induction of articular-like chondrogenesis from MSCs, implying that the paracrine factors supplied are crucial to the generation of functionally superior neocartilage.

**Figure 6:**
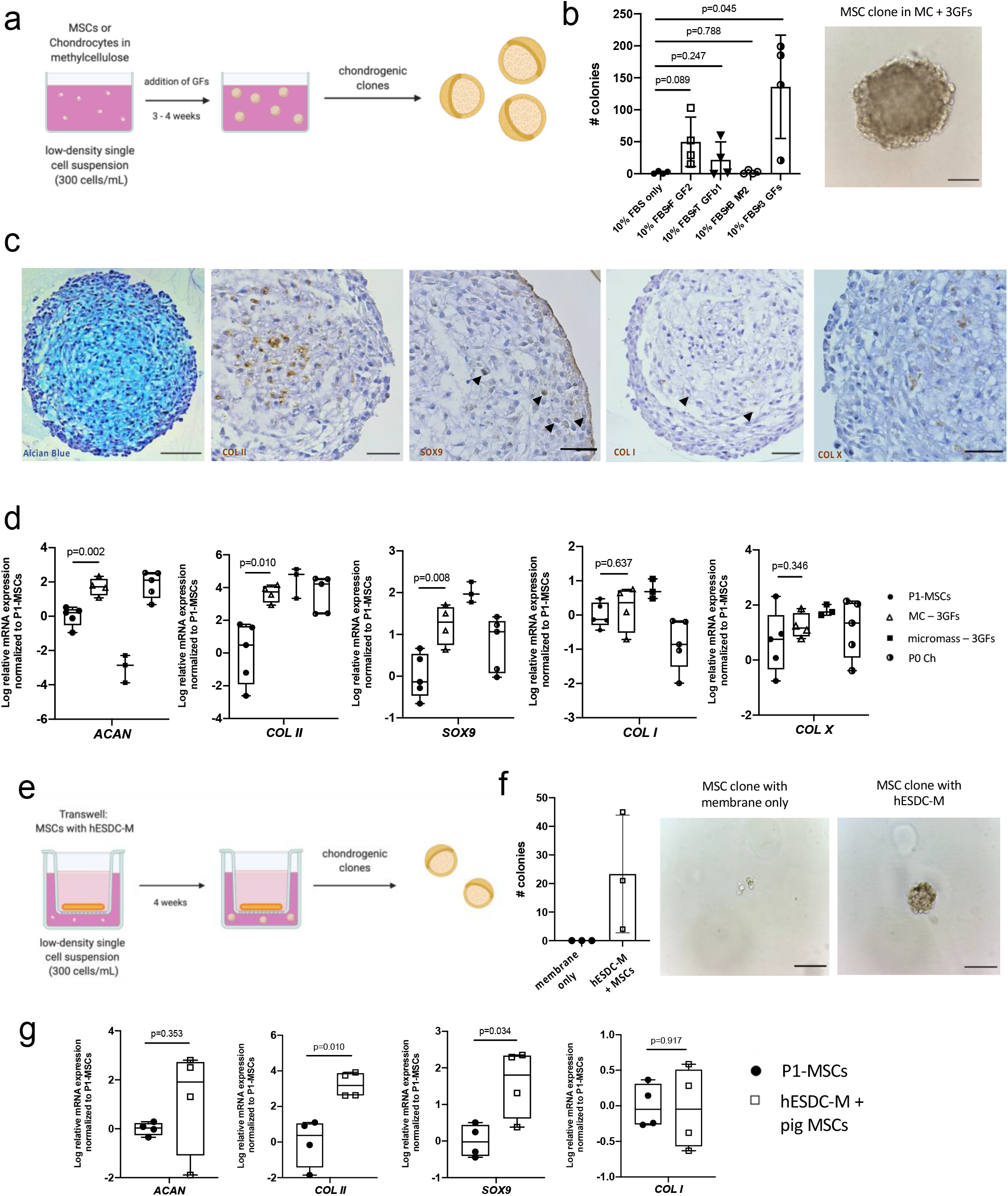
hESDC-M produce paracrine factors that drive chondrogenesis of endogenous cells. (a) Schematic depicting the methylcellulose (MC) culture method. (b) Clonogenicity of bone-marrow derived mesenchymal stromal cells (MSCs) in MC with different GFs; n=4 biological replicates. Representative image of a pig MSC-derived colony after 4 weeks in MC with 3 growth factors (right); scale bar = 100 μm. (c) Alcian Blue staining (left) and immunohistochemical staining various chondrogenic markers of pig MSCs grown in MC with 3 GFs after 4 weeks. Scale bar = 100 μm. (d) qPCR of chondrogenic genes (n=5 biological replicates for P1-MSCs and P0 Ch, n=4 for MSCs in MC, and n=3 for MSCs cultured micromass). (e) Schematic of the MC with Transwell culture method. (f) Clonogenicity of pig MSCs in MC with a membrane only or hESDC-M in Transwell after 4 weeks, n=3 biological replicates per group. Representative images of MSCs in the Transwell after 4 weeks are shown; scale bar = 100 μm. (g) qPCR of chondrogenic genes from pig MSCs grown in Transwell with hESDC-M, n=4 biological replicates. p-values were calculated with an unpaired Student’s t-test; data presented as mean ± SD or box and whisker plots showing all points.

## Discussion

We have shown that hESC-derived chondrocytes administered as a cryopreservable, membrane-embedded formulation support clinically relevant, long-term repair of full-thickness articular cartilage defects in pigs. At the molecular level, membranes were found to contain no detectable residual hESCs and be populated with immature chondrocytes resembling both embryonic chondroprogenitors and fetal juvenile articular chondrocytes. Based on the high expression levels of *SOX9* and genes associated with proliferation, it is likely that these immature articular chondrocytes mature *in vivo* to upregulate matrix production and assume a proper zonal identity^18^. Critically, the hyaline-like cartilage found in repaired defects had similar biomechanical properties to naïve tissue, further supporting the concept that hESDC-M can adopt adult-like properties upon transplantation and integration with surrounding cells and/or support recruitment and differentiation of chondrogenic cells into hyaline cartilage via paracrine factors.

Despite being a xenograft in immunocompetent recipients with no immunosuppression, we found no evidence of local inflammation or immune cell infiltration. In light of the clinical use of particulate juvenile chondrocytes and osteochondral transplants as allograft material^38,39^, this is not entirely unexpected. However, other studies have demonstrated that xenografted articular chondrocytes in the knee can elicit a severe immune response^40^. Once transitioned to human trials of allogenic hESDC-M for focal articular cartilage repair, patients will have to be evaluated carefully to determine which of them may require additional screening or exclusion based upon the degree of joint inflammation or other systemic diseases. Additionally, although we did not detect any peripheral dissemination of human cells or residual pluripotent cells in the current work using highly sensitive analyses, each of these will require additional testing to validate the safety profile of this potential cell therapy and the release criteria for each production batch.

The significant biomechanical improvement seen in defects treated with human cells is very likely the result of both autocrine and paracrine mechanisms. This suggests that factors secreted by hESDC-M such as BMP-2 can recruit endogenous cells capable of producing hyaline-like cartilage and support their differentiation down this path. Murphy et al. recently demonstrated that implantation of hydrogels loaded with BMP-2 and sVEGFR-1 could significantly improve the outcome of microfracture in both young and old mice^9^. Moreover, they show that treatment with BMP-2 alone was insufficient to generate articular cartilage following microfracture, clearly indicating that induction of an articular-like fate from MSCs requires more than one input. Consistent with other studies^41,42^ our data show that BMP-2 was not secreted at detectable levels by hBM-MSCs but was robustly generated by hESDC-M; moreover, the scRNA-Seq data presented here clearly identify a population similar to primitive, chondroinductive cells present in the developing limb. In addition to chondrogenic factors, it is possible that pro-inflammatory signaling needs to be blunted to allow for optimal regeneration by endogenous cells. Following the exposure of the knee joint and creation of the defects, significant local inflammation occurs, driving production of catabolic enzymes and promoting fibrocartilage formation which may be ameliorated via the paracrine production of counteracting proteins such as TIMP-1/2 and IL-1RA (a decoy receptor for the pro-inflammatory cytokine IL-1α^43^) by hESDC-M. TIMP-1 and -2 are inhibitors of matrix metalloproteinases and have shown to be chondroprotective^44,45^ and prevent matrix loss downstream of the pro-inflammatory cytokine IL-1β^46^. Together, these potential chondrogenic paracrine factors may modulate the microenvironment of the defect to promote migration and differentiation of endogenous cells into articular cartilage. It will be important to assess the response when hESDC-M are used in injuries that occurred significantly before transplantation, as acute inflammation will have subsided and chronic inflammation may be elevated.

There are caveats to the data presented here. Although pig models of cartilage repair are commonly used^23,24^, some biomechanical differences between human and pig joints are well documented^25^. Porcine knees have significantly softer cartilage and longer trochlea than those of humans, putting the joint in a constant state of flexion. Moreover, humans are more active than pigs, leading to higher load bearing properties on average. These parameters could influence implanted cells in ways different than documented here. In addition, focal articular cartilage defects in humans often involve the cartilage without reaching the subchondral space^24^ and are often latent for some time prior to a repair procedure while in the current study, focal full-thickness lesions were freshly made; in addition, we did not address the impact on subchondral bone via microCT although general histological analysis of the subchondral bone was conducted. Many model species including large animals can spontaneously heal partial-thickness defects without bone marrow stimulation^23,24^, highlighting the lack of a perfect pre-clinical model for articular cartilage repair. In line with this, integration of graft tissue with endogenous articular cartilage is a major concern for cell-based therapies of full-thickness defects^14,23^ as transplanted autologous chondrocytes do not appear to consistently attract cells with capacity to promote bridging; graft integration failure has been documented in a pig model of marrow stimulation as well^23^. We did not include control groups in the current study that tested transplantation of autologous chondrocytes, MSCs or allogeneic hBM-MSCs as these cells have been vigorously evaluated in past studies^23,47,48^, and have not demonstrated long-term retention of implanted cells. Based on the relatively primitive nature of hES-derived chondrocytes, we speculate their increased motility, secretome, and integration will support better graft integration outcomes in full-thickness defects. Finally, pain and the associated loss of mobility are of significant concern in patients with focal cartilage lesions^14^. Although we did not directly assess joint usage or gait as a surrogate for pain in this study, future clinical evaluations of hESDC-M at all stages will include assessments of pain with reproducible test re-test results^49^.

In spite of these potential weaknesses, the presented robust potential of hESDC-M to enact articular cartilage repair as a universal, off-the-shelf product should be explored clinically. Current cell therapies for focal defect repair rely on expansion and implantation of autologous cells (e.g. MACI) or allogenic juvenile articular cartilage which has limited availability. If hESDC-M can enact meaningful repair of focal defects in patients, this represents a unique opportunity to scale an allogenic therapy, providing the possibility of superior functional repair from an inexhaustible source of chondrogenic cells.

## Acknowledgments

Schematic for *in vitro* studies (Figure 6) were created with Biorender.com. De-identified human cartilage samples were collected under UCLA IRB# 10-001857. Research reported in this publication was supported by the National Institute of Arthritis and Musculoskeletal and Skin Diseases of the National Institutes of Health under Awards K01AR061415 and R01AR071734 to DE and the National Institute of Aging Award R01AG058624. The content is solely the responsibility of the authors and does not necessarily represent the official views of the National Institutes of Health. This work was also supported by NIH grants DOD grant W81XWH-13-1-0465 and CIRM grants RB5-07230 and TRAN1-09288, all to DE; and NIH R35 DE027550 grant to GDC.

---

**Figure S1:**
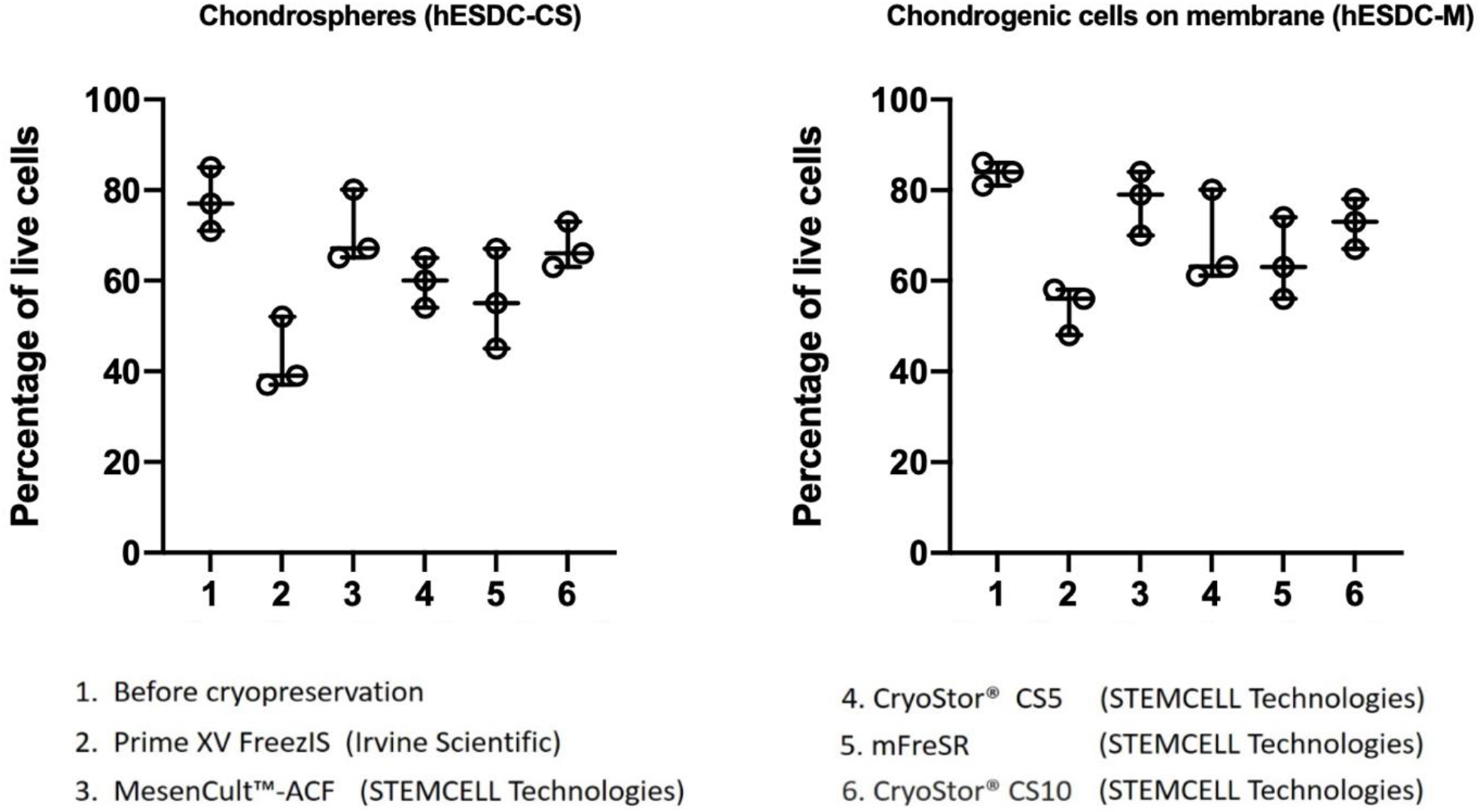
Optimization of cell cryopreservation. Cell viability was assessed using Live/Dead assay prior to viably freezing and within 4 hours after thawing. Mesencult™ ACF provided the best viability post-thaw and was used for all subsequent preparations. Data are presented as mean ± SD of 4 biological replicates.

**Figure S2:**
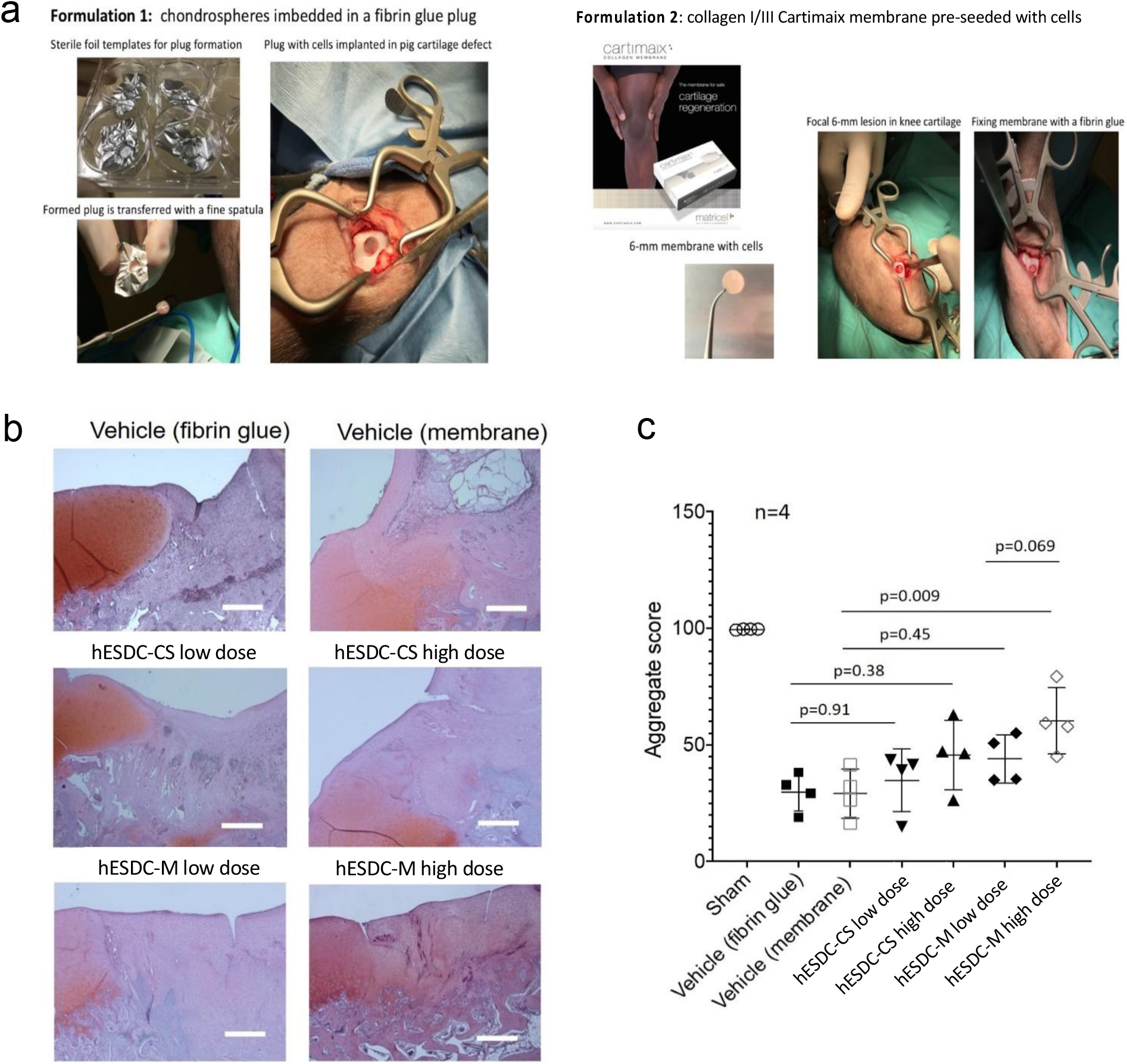
Overview of surgical procedure and assessment of short-term cartilage repair at 1 month. (a) Full-thickness defects were created in Yucatan minipig knees and transplanted with low and high doses of either chondrospheres (CS) or hESDC-M. (b) Assessment of proteoglycan content via Safranin O staining one month after implantation; images show the junction of intact articular cartilage with the defect area. (c) ICRS II scoring of defects demonstrated the high dose, membrane embedded formulation of hES-derived chondrocytes provided superior short-term repair. n=4 defects per condition, 2 defects per knee. Scale bars = 100 μm; data presented as mean ± SD. p-values were calculated via one-way ANOVA followed by Tukey’s test.

**Figure S3:**
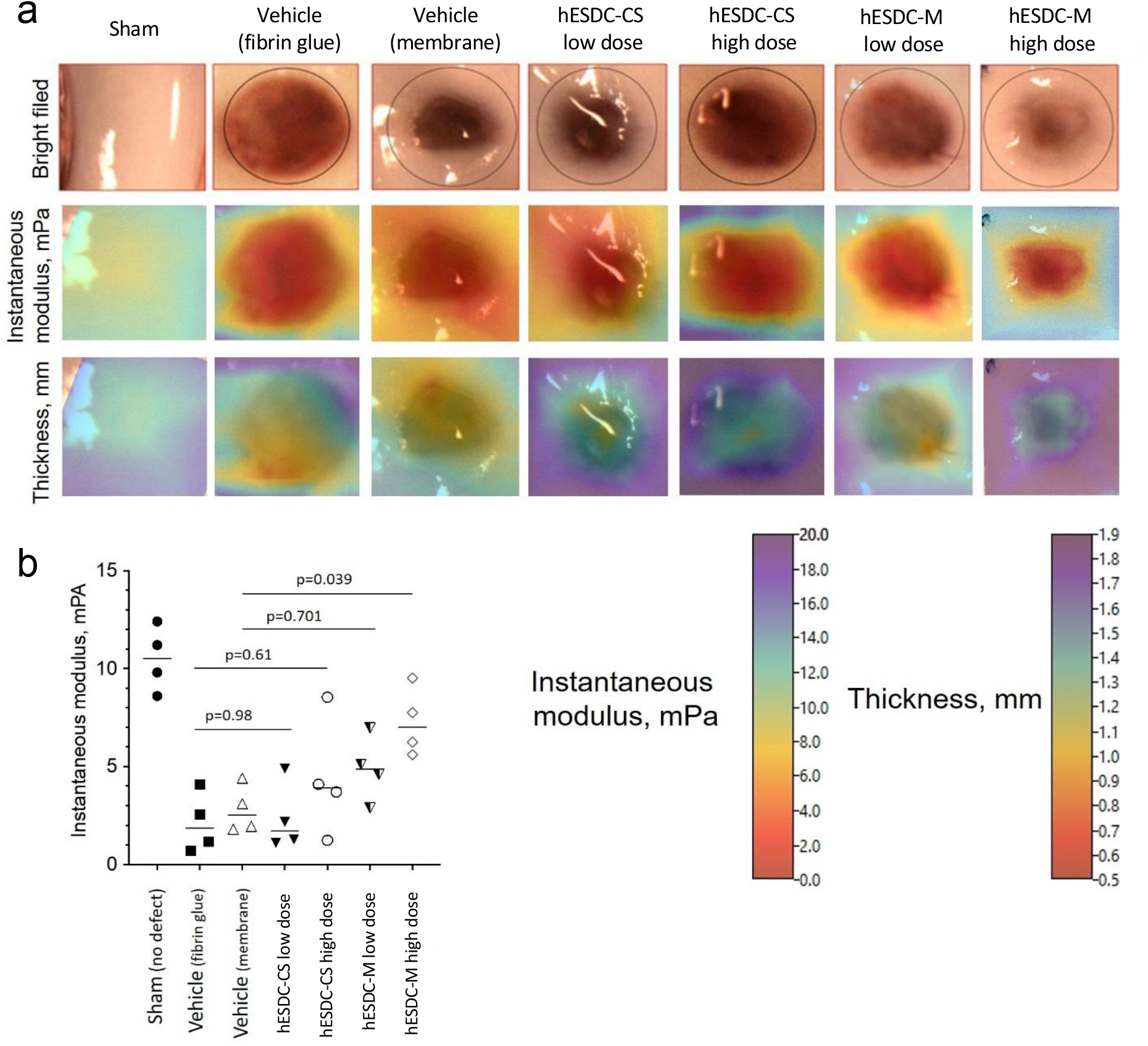
Biomechanical properties of repaired cartilage defects. (a) Analysis of biomechanical properties was carried out using Mach-1 scanning indenter (Biomomentum, Canada). At least 10 points were analyzed within each defect. Heat maps representing instantaneous modulus and thickness are shown for each group. Color mapping is artificial to illustrate the differences between specimens; scales for each measurement are shown below. (b) Quantitative assessment of instantaneous modulus within the healing defect area 1 month after transplantation of 2 doses of either hESC-derived chondrospheres (hESDC-CS) or collagen I/III membrane embedded hESDC-M. The best repair was observed in the high dose hESDC-M group. Data presented as mean ± SD of aggregate values for 4 defects per each group; p-values were calculated via one-way ANOVA followed by Tukey’s test.

**Figure S4:**
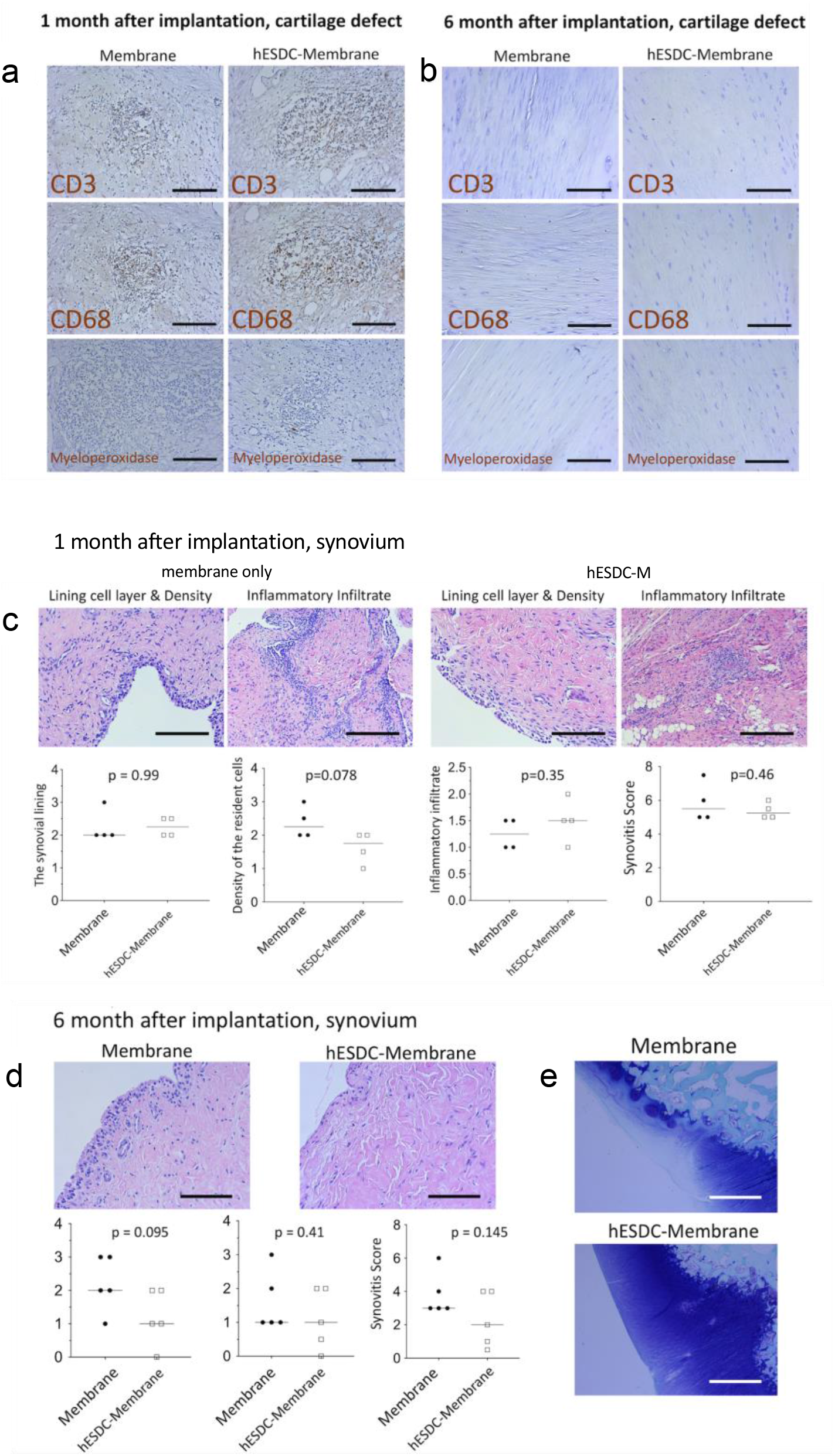
Immunological response to hESDC-M. (a) Representative images of immunohistochemical staining for immune cell markers in the cartilage defect at 1 month post implantation of the membrane only (left column) or the hESDC-M (right column). (b) Representative images of immunohistochemical staining for immune cell markers in the cartilage defect at 6 months post implantation of the membrane only (left column) or the hESDC-M (right column). (c) Representative Hematoxylin and Eosin (H&E) stains and quantification of synovial characteristics 1 month after implantation using previously described methods^29^ (n=4 biological replicates for membrane only and hESDC-M group (d) Representative H&E stain and corresponding quantification of synovium 6 months after implantation; left image is the membrane only, right image is the defect with hESDC-M; (n=5 biological replicates for membrane only group, n=5 for hESDC-M group) (e) Representative Toluidine Blue stain of articular cartilage with the defect area 6 months after implantation; top image is the membrane only, bottom image is the defect with hESDC-M. All scale bars = 100 μm, p-values calculated with an unpaired t-test and data is presented as mean ± SD.

**Figure S5.**
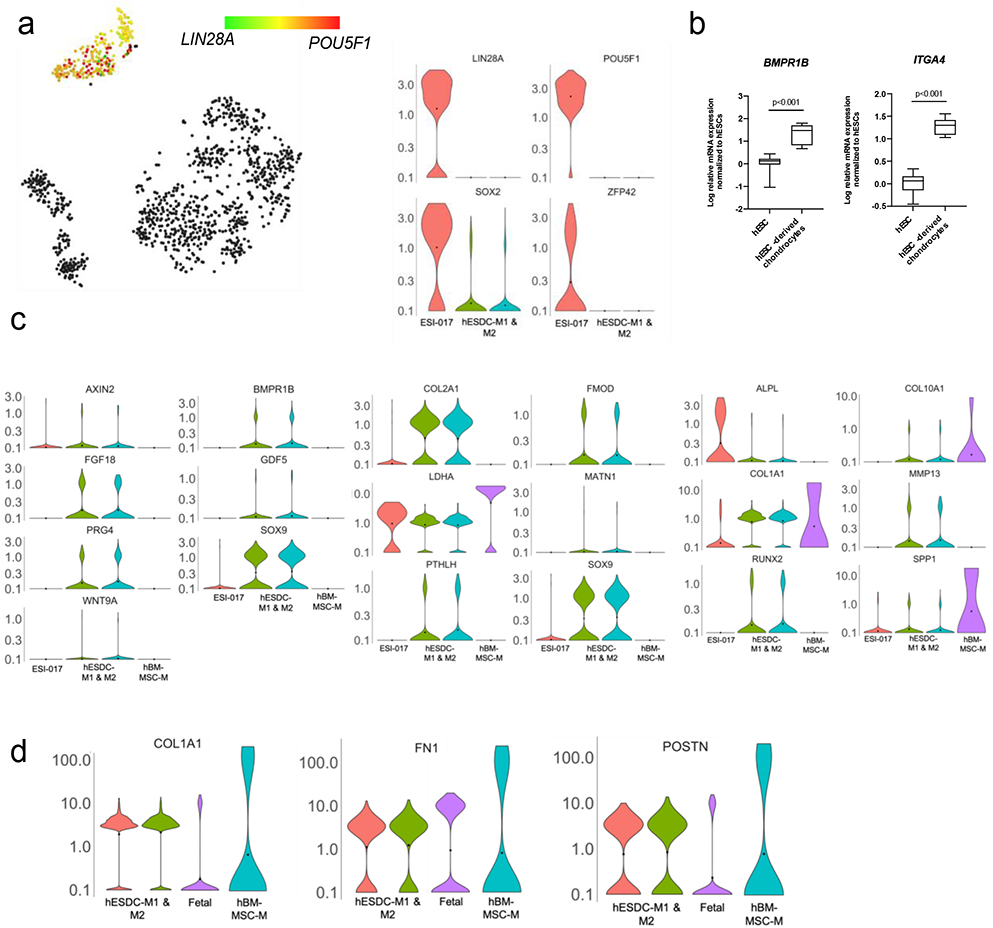
hESDC-M are not contaminated by hESCs and represent more immature chondrogenic cells than hBM-MSC-M. (a) t-SNE and violin plots depicting expression of indicated genes at single cell resolution. (b) qPCR of superficial gene expression (n=7 independent batches; p-values were calculated with an unpaired t-test; data presented as box and whisker plots showing all points. (c) Violin plots for gene expression of selected chondrogenic genes in hESCs, hESDC-M and human bone marrow MSCs cultured on membranes (hBM-MSCs-M). (d) Violin plots for gene expression of selected mesenchymal genes.

**Figure S6.**
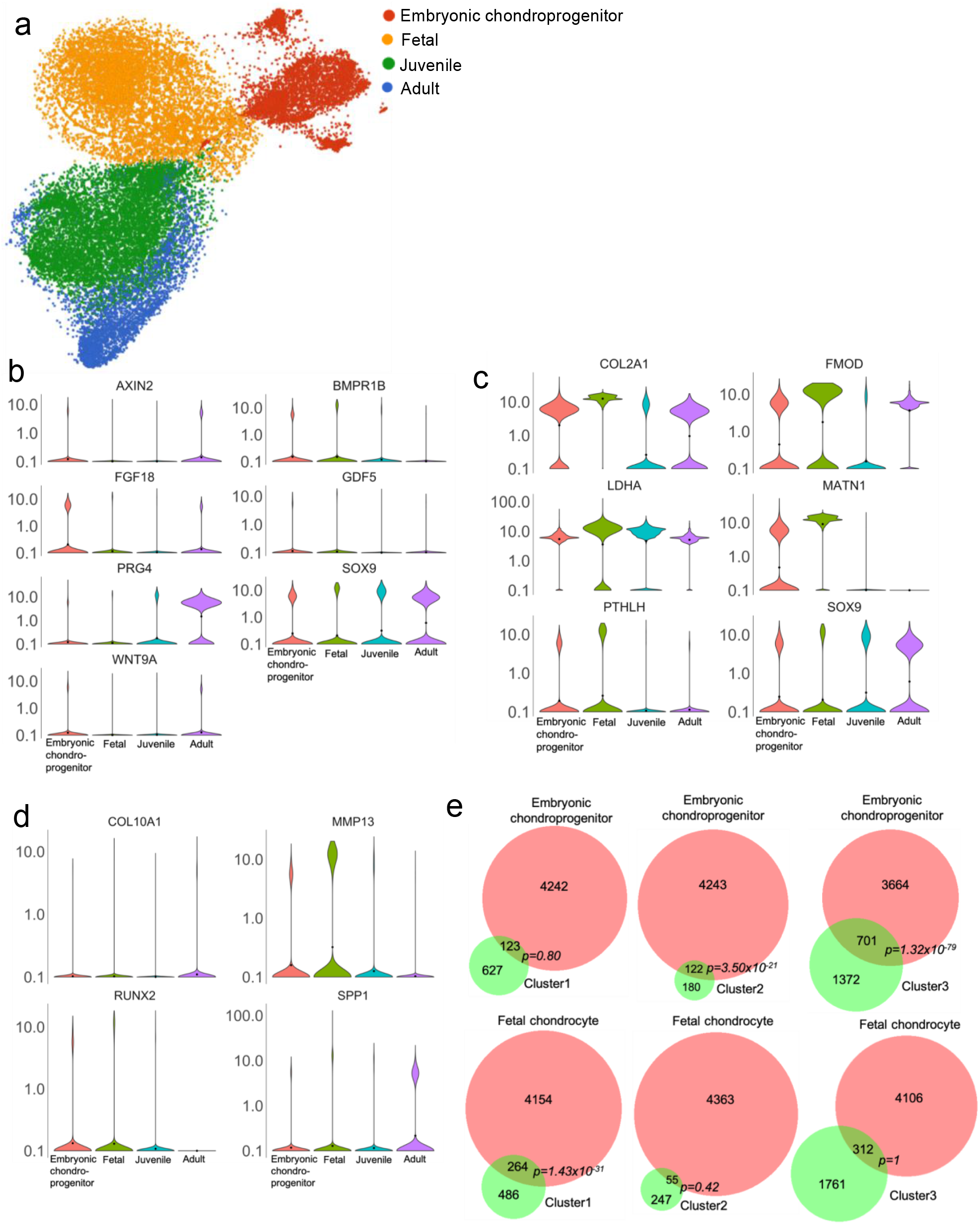
Definition of chondrogenic ontogeny at the single cell level. (a) t-SNE plot of single cell sequencing data generated at 4 stages of human chondrocyte ontogeny. Violin plots for gene expression of selected (b) superficial, (c) transitional and (d) deep zone genes. (e) Gene sets were created by intersecting two previously published data sets^17,18^ generated with bulk RNA-seq of embryonic chondroprogenitors and fetal chondrocytes. Genes enriched in “pre-chondrocytes” vs. “resting chondrocytes” were overlapped with genes enriched in “embryonic 5-6 WPC (weeks post conception)” and “17 WPC” and vice versa to create lists of common genes enriched in embryonic chondroprogenitors and fetal chondrocytes. Venn diagrams demonstrating overlap of biomarker genes of the indicated cluster of hESDC-M (see Figure 2h) analyzed by scRNA-Seq with gene lists generated by bulk RNA-Seq.

**Figure S7:**
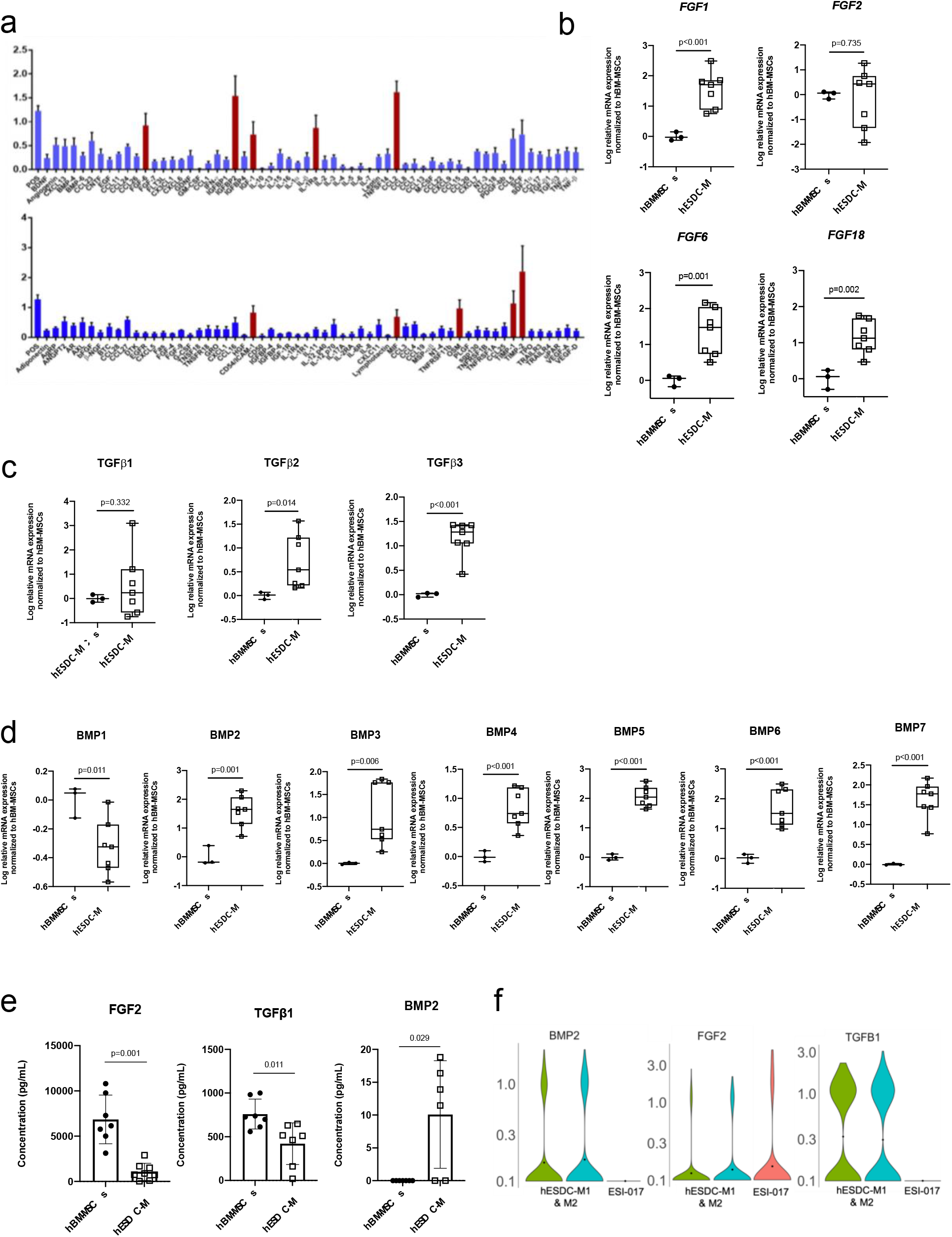
Paracrine factor analysis of hESDC-M. (a) Lysates from hESDC-M were assessed for the presence of soluble proteins/cytokines (n=3 batches). Red bars indicate factors secreted at levels enriched compared to other proteins. Data presented as mean ± SEM of 3 biological replicates. qPCR for various (b) FGF, (c) TGF-**β** and (d) BMP family members in P1 hBM-MSCs (n=3 biological replicates of 27-29 yo) and hESDC-M (n=7 batches). (e) ELISA analyses of representative growth factors (BMP-2, TGF-**β**1, & FGF-2) secreted by P1-3 hBM-MSCs (n=7 biological replicates aged 19-64 yo) and hESDC-M (n=9 batches for FGF2, n=7 batches for TGF-**β**1 and n=6 batches for BMP-2). p-values calculated with unpaired t-test; error bars represented as mean ± SD. (f) Violin plots depicting gene expression in ESI-017 cells and 2 replicates of hESDC-M analyzed by scRNA-Seq.

**Figure S8:**
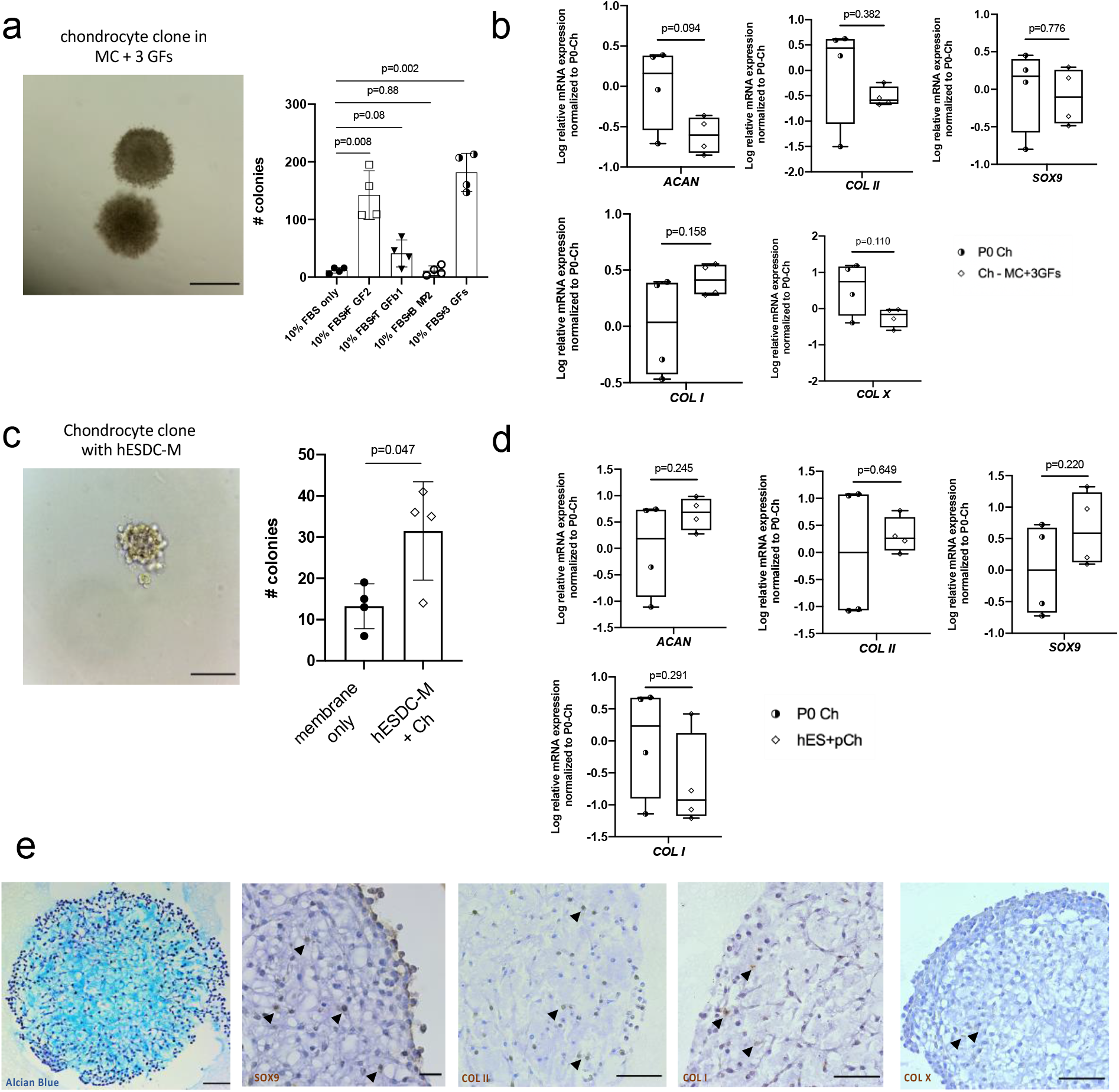
Endogenous articular chondrocytes maintain their chondrogenic profile in the presence of hESDC-M. (a) Representative image of two pig chondrocyte (Ch) colonies (scale bar = 500 µm) & clonogenicity of pig chondrocytes (n=4 biological replicates) cultured in methylcellulose (MC) for 4 weeks. (b) qPCR of chondrogenic genes (n=4 biological replicates). (c) Representative image of a pig Ch colony in MC after 4 weeks of culture with hESDC-M in a Transwell. Clonogenicity of pig Ch in MC with either an empty membrane or hESDC-M in a Transwell after 4 weeks (n=4 biological replicates per group). Scale bar = 100 µm. (d) qPCR of chondrogenic genes in pig Ch grown in Transwell culture with hESDC-M (n=4 biological replicates). (e) Alcian Blue staining (left) and immunohistochemical staining of various chondrogenic markers of pig chondrocytes grown in MC with 3 GFs after 4 weeks. Scale bar = 100 μm; all p-values calculated with unpaired t-test; error bars represented as mean ± SD.

