## Supplemental methods for "Long-term repair of porcine articular cartilage using cryopreservable, clinically compatible human embryonic stem cell-derived chondrocytes"

**General Methods:** For all experiments, biological replicates were employed to generate data. For *in vitro* experiments expected to yield large differences, standard practice of using 3 replicates was followed. All statistical methods are described in the figure legends.

**Culture of pluripotent stem cells and generation of skeletal progenitors:** For all experiments, research grade ESI-017 were used; the stock line was purchased from BioTime. Cells were cultured on ESC-grade Matrigel (Corning) in mTesR1/Plus media and passaged using ReLeSR (Stemcell Technologies) according to manufacturer's instructions. Batch scale production of ESI-017 cells was carried out by seeding ~5 million cells in CellSTACK-5 flasks, yielding ~1 billion cells per batch. Each batch was tested for mycoplasma contamination by PCR (Sigma). Cells were then transferred to suspension culture in mTesR1/Plus for up to 5 days in spinner flasks (60 RPM) for additional expansion and differentiation. Differentiation was conducted as described previously[17,18,27]. Briefly, mTesR1/Plus was replaced with Mesoderm Induction Media A (MIM-A) for 3 days [X-Vivo (Lonza) containing ROCK inhibitor (10 μM; Y27623, Tocris), FGF2 (10ng/ml), Wnt3a (10ng/ml) and Activin A (10ng/ml)] followed by MIM-B for 3 days [X-Vivo containing Wnt3a (10ng/ml), Noggin (50ng/ml) and FGF-2 (10ng/ml)]. MIM-B contained no ROCK inhibitor or Activin A to avoid excessive endodermal differentiation. After 7 days of differentiation, when skeletal progenitors were already specified, media was changed to Chondrogenic Induction Media A (CIM-A; X-Vivo containing BMP-4 (10ng/ml) and FGF-2 (10ng/ml)) for 4 additional days. For 2 hours before starting MACS isolation, ROCK inhibitor was added again to the cell culture media to increase cell survival after enzymatic dissociation and sorting, and these skeletal progenitors were then isolated using MACS by depleting for CD326 (PerCP) and CD309 (PE), yielding ~0.6 billion cells; or 15-20% of total cells. The remaining cells represented undifferentiated cells, endodermal precursors, and also mesodermal cells committed to cardiovascular and hematoendothelial lineages[27]. The purity of cells isolated from MACS was routinely assessed for negativity of CD326 (PerCP) and CD309 (PE) using flow cytometry.

**Generation of chondrospheres (CS):** Skeletal progenitors from MACS were cultured in EZSphere plates (Nacalai USA) at 100,000 cells per well in CIM-B[17,18] (X-Vivo containing Shh (25ng/ml), ROCK inhibitor (10µM), BMP-4 (50ng/ml), FGF-2 (10ng/ml), IGF-1 (10ng/ml) and Primocin (100µg/ml)) in 5% oxygen for 3-5 days to form chondrogenic aggregates. After firm aggregates were verified with microscopy, they were transferred to a perfusion bioreactor (Applikon Biotech) in Maturation Media (MM; X-Vivo media containing FGF-2 (10ng/ml), BMP-4 (1ng/ml), IGF-1 (10ng/ml), LIF (50ng/ml), TGF-β1 (10ng/ml) and Primocin (100 µg/ml)) until d40 of differentiation. Conditions in the bioreactor were kept at 37°C, 5% $O_2$, 5% $CO_2$ and 30 RPM. Fresh media was added at a rate of 1 mL/hour. Upon maturation, chondrospheres contained 5 x $10^4$ cells on average. At d40, chondrospheres were cryopreserved in Mesencult-ACF plus ROCK inhibitor (10 µM) and stored in liquid nitrogen until use.

**Generation of hES-derived chondrocytes on membranes (hESDC-M):** Skeletal progenitors isolated via MACS were seeded onto porcine collagen I/III membranes (Cartimaix; Matricel) sized with a 6mm biopsy punch at two different densities (1 or 4 million viable cells), yielding ~6 x $10^5$ or 3 x $10^6$ cells attached to the membranes, respectively. Membranes were then cultured in 5% oxygen for 3-5 days in CIM-B and then transferred to the bioreactor as above for chondrospheres with the exception that rotation was not started until 3 days after transfer of membranes to allow the cells to attach to the membranes. At d40, membranes were cryopreserved using Mesencult-ACF as above for chondrospheres.

**Generation of hBM-MSCs on membranes (hBM-MSC-M):** Human bone marrow-derived mesenchymal stromal cells (n=3 donors aged 19, 68, and 87 years (all P1);, pooled) were seeded onto porcine collagen I/III membranes (Cartimaix; Matricel) sized with a 6 mm biopsy punch at a density of $3 \times 10^6$. Membranes were then cultured in 5% oxygen for 3-5 days in CIM-B, then maintained with MM until d40. At d40, membranes were digested for single cells and scRNA-sequencing was performed.

**Optimization of cryopreservation media:** Live cell numbers per batch of chondrospheres or membranes were determined prior to freezing using the Live/Dead Cell Viability Assay (Biovision). Chondrospheres or membranes were then cryopreserved following the manufacturer's instructions for each product (Prime XV FreezIS, Irvine Scientific; Mesencult-ACF, CryoStor CS5 or 10, mFreSR; Stemcell Technologies). Viability post-thawing from the same batch was compared to the starting viability before freezing.

**Quantitative Real-Time PCR:** Power SYBR Green (Applied Biosystems) RT-PCR amplification and detection was performed using an Applied Biosystems Step One Plus Real-Time PCR machine. The comparative Ct method for relative quantification (2-ΔΔCt) was used to quantitate gene expression, and displayed as $Log_{10}$ of relative expression. TBP (TATA-box binding protein) or RPL7 (ribosomal protein L7) was used for gene normalization. For quantification of human cell numbers in pig samples, a human *TERT* Taqman assay was used (Thermo). A standard curve was created with known numbers of human cells, which both determined the detection threshold as well as allowed calculation of human cell numbers in a sample based on Ct values.

**Primer sequence List:** Please see Table 1 for a list of qPCR primer sequences used

**Table 1**

| Gene name | Primer Sequence | GenBank Accession |
|---|---|---|
| hRPL7 | Forward: 5’ CCAAATTGGCGTTTGTCATCAG 3’<br>Reverse: 5’ GCATGTTAATCGAAGCCTTGTTG 3’ | NM_000971 |
| hTBP | Forward: 5’ TGCACAGGAGCCAAGAGTGAA 3’<br>Reverse: 5’ CACATCACAGCTCCCCACCA 3’ | NM_001172085 |
| hCOL2A1 | Forward: 5’ TGGACGATCAGGCGAAACC 3’<br>Reverse: 5’ GCTGCGGATGCTCTCAATCT 3’ | NM_001844 |
| hSOX9 | Forward: 5’ AGCGAACGCACATCAAGAC 3’<br>Reverse: 5’ GCTGTAGTGTGGGAGGTTGAA 3’ | NM_000346 |
| hLIN28b | Forward: 5’ CATCTCCATGATAAACCGAGAGG 3’<br>Reverse: 5’ GTTACCCGTATTGACTCAAGGC 3’ | NM_001004317 |
| hPOU5F1 | Forward: 5’ AGTGAGAGGCAACCTGGAGA 3’<br>Reverse: 5’ CACTCGGACCACATCCTTCT 3’ | LC006945.1 |
| pRPL7 | Forward: 5’ CAGGATCAGAGGTATCAA 3’<br>Reverse: 5’ TATATGGTTCCACAATTCTC 3’ | NM_001113217.1 |
| pACAN | Forward: 5’ CTACGACGCCATCTGCTACA 3’<br>Reverse: 5’ CTTCACCCTCGGTGATGTTT 3’ | NM_001164652.1 |
| pCOL2A1 | Forward: 5’ GAGAGGTCTTCCTGGCAAAG 3’<br>Reverse: 5’ AAGTCCCTGGAAGCCAGAT 3’ | XM_021092611.1 |
| pSOX9 | Forward: 5’ CCACCGAAGAAAGACCGTAA 3’<br>Reverse: 5’ CTTGGAATGTGGGTTCGAGT 3’ | NM_213843.2 |
| pCOL1A1 | Forward: 5’ CCAGTCACCTGCGTACAGAA 3’<br>Reverse: 5’ ACGTCATCGCACAACACATT 3’ | LC223106.1 |
| pCOLXA1 | Forward: 5’ ACTTCTCCTACCACATTC 3’<br>Reverse: 5’ CCATACCTGGTCATTATCT 3’ | NM_001005153.1 |

h = human, p = pig

**Large animal model of articular cartilage repair**

Yucatan minipigs were purchased from S & S farms at 6 months of age and housed under the supervision of the USC Department of Animal Resources (DAR). All pre-operative, surgical and post-operative procedures were conducted following USC DAR guidelines and were overseen by the USC Institutional Animal Care and Use Committee (IACUC). Five animals per group (main study) with 2 defects each were included based on power calculations to yield the minimum number of animals projected to acquire statistically significant results. Briefly, we used the formula $n = 1 + 2C\left(\frac{s}{d}\right)^2$ based on a 30-40% difference and 20% standard deviation between experimental groups, with these parameters based on our previous studies of cell-based focal repair of articular cartilage in large animal models[47-50]. Animals were anesthetized for surgery using Telazol/Xylazine 2.2-4.4 mg/kg administered intramuscularly. Medial para-patellar arthrotomy was performed by inserting microsurgical scalpel medially and proximally to the insertion of the patellar tendon on the tibia and extending it proximally until the attachment of the quadriceps muscle. The medial margin of the quadriceps was separated from the muscles of the medial compartment. The joint was extended and the patella dislocated laterally. The joint was then be fully flexed to expose the patellar groove. A 6 mm-diameter disposable biopsy punch was used to create two full-thickness injuries in the articular cartilage, without perturbation of the underlying subchondral bone. The wound bed was cleaned with sterile cotton applicators. 100 $\mu$L of fibrin glue (Ethicon) was used to suspend the chondrospheres in a sterile formed plug for implantation. The formed chondrosphere plug was then transferred and compressed into the defect. Both the chondrospheres and hES-derived chondrocyte membranes were sealed with several droplets of fibrin glue to allow setting of the implantation. No randomization was applied. For animals receiving chondrospheres in the pilot study, surgical fibrin glue with or without chondrospheres was applied directly to the defect areas. For animals receiving membranes, pre-sized membranes were applied to the defect and secured with fibrin glue. Arthrotomies were then relieved, wounds sutured and animals monitored for recovery. Following 1 or 6 months, animals were euthanized after receiving anesthesia as above using pentobarbitol (100 mg/kg).

**Tissue collection and digestion:** Adult human primary tissue samples were obtained from National Disease Research Interchange (NDRI). Fetal and embryonic tissue samples were obtained from Novogenix Laboratories. All donated material was anonymous, carried no personal identifiers and was obtained after informed consent. Sex of the specimens was unknown. Human primary tissues, hES-derived chondrocytes on membranes, or hBM-MSCs on membranes were manually cut into small pieces and digested 4-16hrs at 37° C with mild agitation in digestion media consisting of DMEM/F12 (Corning) with 10% FBS (Corning), 1 mg/mL dispase (Gibco), 1 mg/mL type 2 collagenase (Worthington), 10 µg/mL gentamycin (Teknova) and 100µg/ml primocin (Invivogen).

**Histology:** Tissues were fixed in 10% formalin and sectioned at 5 µm[18]. For DAB immunohistochemical (IHC) staining, sections were deparaffinized using standard procedures and antigen retrieval was performed by incubating the samples in 1x citrate buffer pH 6.0 (Diagnostic Biosystems) at 60°C for 30 minutes, followed by 15 minutes cooling at room temperature. Endogenous peroxidase activity was quenched by treating samples with 3% $H_2O_2$ for 10 minutes at RT. Sections were then blocked in 2.5% normal horse serum for 20 minutes. Sections were then incubated with primary antibodies diluted in TBS with 0.1 - 1% BSA (Sigma) overnight at 4° C. Sections were washed 3 times with TBS + 0.05% Tween 20 (TBST, Sigma) before addition of HRP-conjugated secondary antibody for a 30-minute incubation at RT. Sections were washed 3 times with TBST after secondary incubation and DAB substrate was then added until positive signal was observed. Sections were then immediately washed with tap water, counterstained in hematoxylin for 15 - 30 seconds and washed again with tap water before dehydration and mounting. Isotype controls or secondary antibody only controls were used for IHC. For Hematoxylin and Eosin staining, sections were deparaffinized, rinsed in tap water, and stained with Hematoxylin for 3 minutes. Sections were then washed in tap water and stained with Eosin for 2 minutes before a final wash in tap water. Safranin O/Fast Green staining was performed as previously described[51,52]. To quantitate cartilage repair, the ICRS II scoring system[33] was employed by two blinded observers. Toluidine Blue and Alcian Blue staining was performed on deparaffinized sections in accordance with standard laboratory techniques.

**Biomechanical assessment of defect repair:** Freshly harvested cartilage tissues were affixed to the sample holder of the Mach-1 Mechanical Tester (Biomomentum) using instant glue (Loctite 4013) and immersed in DMEM/0.9% NaCl (1:1). The Mach-1 configuration uses a spherical indenter tool attached to a highly sensitive multiaxial load cell and automated fine motor controller, allowing for compression of the cartilage by 30% of its thickness, and live recording of resultant forces generated in all x-y-z planes. The indenter tool is then replaced with a needle which penetrates the cartilage at each point until forces comparable to underlying bone are detected, allowing for accurate thickness measurement. Altogether, the forces generated during indentation and thickness measurements are used to calculate the instantaneous modulus, which reflects the elasticity, stiffness, and resistance to compression of the tissue. Each condyle was manually mapped for testing using the Biomomentum mapping software. On average, between 40-50 points were tested on each affected condyle, with no less than 10 points directly in the defect areas, and usually about 1-3mm apart from each other on the surface. Each control condyle was tested for an average of 15-20 points, which were spaced about 2-5 mm apart. Mapping coordinates were input into the software and indentation analysis to map instantaneous modulus was performed with the 1 mm spherical indentor tool with the following parameters: Z-contact velocity: 0.1000 mm/s; contact criteria: 0.1003 N; scanning grid: 0.2000 mm; indentation amplitude: 0.300 mm; indentation velocity: 0.300 mm/s; relaxation time: 5s. To map thickness, the indentor tool was replaced with a 26G ¾ inch hypodermic needle and the mapping executed with the following needle penetration parameters: stage axis: position z; load cell axis: Fz; direction: positive; stage velocity: 0.2000 mm/s; contact criteria: 2.000 N; stage limit: 15 mm; stage repositioning: 2x load resolution; offset: 0. Heat maps were generated with the Mach-1 software provided by Biomomentum.

**Single-cell sequencing using 10X Genomics:** Single cell samples were prepared using Single Cell 3′ Library & Gel Bead Kit v2 and Chip Kit (10X Genomics) according to the manufacturer's protocol. Briefly samples were FACS sorted using DAPI to select live cells followed by resuspension in 0.04% BSA-PBS. Nearly 1,200 cells/µl were added to each well of the chip with a target cell recovery estimate of 8,000 cells. Thereafter Gel bead-in Emulsions (GEMs) were generated using GemCode Single-Cell Instrument. GEMs were reverse transcribed, droplets were broken and single stranded cDNA was isolated. cDNAs were cleaned up with DynaBeads and amplified. Finally, cDNAs were ligated with adapters, post-ligation products were amplified, cleaned up with SPRIselect. Purified libraries were submitted to UCLA Technology Center for Genomics & Bioinformatics for quality check and sequencing. The quality and concentration of the purified libraries were evaluated by High Sensitivity D5000 DNA chip (Agilent) and sequencing was performed on NextSeq500.

**10X sequencing data analysis:** Raw sequencing reads were processed using Partek Flow Analysis Software (build version 10.0.21.0210). Briefly, raw reads were checked for their quality, trimmed and reads with an average base quality score per position >30 were considered for alignment. Trimmed reads were aligned to the human genome version hg38-Gencode Genes- release 30 using STAR -2.6.1d with default parameters. Reads with alignment percentage >75% were de-duplicated based on their unique molecular identifiers (UMIs). Reads mapping to the same chromosomal location with duplicate UMIs were removed. Thereafter 'Knee' plot was constructed using the cumulative fraction of reads/UMIs for all barcodes. Barcodes below the cut-off defined by the location of the knee were assigned as true cell barcodes and quantified. Further noise filtration was done by removing cells having >3% mitochondrial counts and total read counts >24,000. Genes not expressed in any cell were also removed as an additional clean-up step. Cleaned up reads were normalized using counts per million (CPM) method followed by log transformation generating count matrices for each sample. Samples were batch corrected on the basis of expressed genes and mitochondrial reads percent. Count matrices were used to visualize and explore the samples in further details by generating tSNE plots generated using default parameters in Partek. K-means clustering was computed for identifying groups of cells with similar expression profile using Euclidean distance metric based on the most appropriate cluster count. A maximum of 1000 iterations were allowed and the top marker features for each cluster was determined.

Gene ontology enrichment analysis for the differentially expressed genes was performed using DAVID Gene Functional Classification Tool (http://david.abcc.ncifcrf.gov; version 6.8). Dot plots and Violin plots were generated in R (v4.0.3) using ggplot2 (v3.3.3) package. Two-way Venn diagrams were generated using BioVenn[53]. Hypergeometric p values were calculated assuming 25,000 human genes.

**Methylcellulose culture method of porcine MSCs and chondrocytes:** porcine bone marrow-derived mesenchymal stromal cells (MSCs) or articular chondrocytes were isolated from the distal femoral epiphysis or articular surface of the condyles, respectively, of 3-4-month-old Yucatan minipigs (S & S Farms). Tissues were digested as described above. Methylcellulose-based media (StemCell) was resuspended with DMEM/F12 (Corning) + 10% FBS + 1% P/S/A and either 10 ng/ml FGF-2, 10 ng/ml BMP-2, 10 ng/ml TGFβ-1, or all three growth factors, to make a 1% methylcellulose-based media. Either P1 MSCs or P0 chondrocytes were seeded in 6-well ultra-low attachment plates (Corning) at a low density (300 cells / ml) and cultured for 3 - 4 weeks. 120 - 160 μL of liquid DMEM/F12 with respective growth factors was applied to the surface of the wells to ensure moisture and nutrients remained available to the cells biweekly. For the Transwell-methylcellulose co-culture, hES-derived chondrocytes on membranes were placed in a 1 μm-pore Transwell insert (Falcon) with DMEM/F12 + 10% FBS + 1% P/S/A. P1 MSCs or P0 chondrocytes were seeded in the same media with methylcellulose, and cultured in

24-well ultra-low attachment plates (Corning) at a low density (300 cells / ml) for 3 – 4 weeks. 40 - 60 µL of media was added biweekly to the methylcellulose as maintenance. Clonogenicity was calculated by manual counting of clones larger than ~40 µm in diameter or greater than 5 cell divisions. All images of clones in methylcellulose were taken on an Echo Revolve Inverted microscope.

**Semi-quantitative detection of proteins secreted by hES-derived chondrocytes on membranes:** Membranes were thawed and washed with serum-free media before being lysed for one hour in RIPA buffer. Lysate was collected and applied to Human Cytokine Antibody Arrays C6 and C7 (RayBiotech) as instructed. Membranes from 3 different batches were used. Spot intensities were measured with Fiji software[54] relative to background and normalized with respect to the average of positive control spot intensities on the same membrane. Normalized values were then averaged across replicates.

**ELISA detection of proteins secreted by hESDC-M:** Either 1 x $10^6$ human bone marrow derived mesenchymal stromal cells (hBM-MSCs; n=7 different donors) or whole hES-derived chondrocytes on membranes (n=6-9 batches) were lysed with 500 µL of 2X Lysis buffer (Ray Biotech) supplemented with a phosphatase/protease inhibitor (Thermo-Fisher). Lysates were then centrifuged to remove cellular debris, and a Bicinchoninic acid (BCA) protein assay (Thermo-Fisher) was performed to quantify total lysed protein. ELISAs for FGF-2 (Ray Biotech), BMP-2 (R & D Systems), and TGF-β1 (R & D Systems) were performed according to the manufacturer’s protocols.

**Antibody List:** Please see Table 2 for a list of antibodies and dilutions used.

**Table 2:** Antibodies used in this study

| **Antibody** | **Vendor** | **Catalog Number** | **Dilution** |
|---|---|---|---|
| CD326-PerCP-Cy5.5 | BD Biosciences | 347199 | 10 uL/$10^6$ cells |
| CD309-PE | R&D Systems | FAB357P | 10 uL/$10^6$ cells |
| Collagen II | Abcam | ab185430 | 1:100 – 1:250 (IHC) |
| PRG4 | Abcam | ab28484 | 1:250 (IHC) |
| SOX9 | Abcam | ab26414 | 1:200 (IHC) |
| Collagen X | Abcam | ab58632 | 1:250 – 1:1000 (IHC) |
| Collagen I | Abcam | ab34710 | 1:250 (IHC) |
| Ku80 | Abcam | ab79391 | 1:250 (IHC) |
| CD3 | Protein Tech | 17617-1-AP | 1:500 (IHC) |
| CD68 | Bioss | BS-1432R | 1:50 (IHC) |
| Myeloperoxidase | Invitrogen | PA5-16672 | 1:50 (IHC) |
| Anti-Mouse IgG ImmPRESS | Vector | MP-7422 | Pre-diluted |
| Anti-Rabbit IgG ImmPRESS | Vector | MP-7401 | Pre-diluted |

**Data availability:** All scRNA-sequencing data are deposited in GEO under accession GSE142045.
